# *Kudoa* genomes from contaminated hosts reveal extensive gene order conservation and rapid sequence evolution

**DOI:** 10.1101/2024.11.01.621499

**Authors:** Claudia C Weber, Michael Paulini, Wellcome Sanger Institute Tree of Life Management, Samples and Laboratory team, Wellcome Sanger Institute Tree of Life Core Informatics team, Mark L Blaxter

## Abstract

Myxozoans are obligate endoparasites that belong to the phylum *Cnidaria*. Compared to their closest free-living relatives, they have evolved highly simplified body plans and reduced genomes. *Kudoa iwatai*, for example, has lost upwards of two thirds of genes thought to have been present in its ancestors. However, little is known about myxozoan genome architecture because of a lack of sufficiently contiguous genome assemblies.

This work presents two new, near-chromosomal *Kudoa* genomes, built entirely from low-coverage long reads from infected fish samples. The results illustrate the potential of using unsupervised learning methods to disentangle sequences from different sources, and facilitate producing genomes from undersampled taxa. Extracting distinct components of chromatin interaction networks allows scaffolds from mixed samples to be assigned to their source genomes. Meanwhile, low-dimensional embeddings of read composition permit targeted assembly of potential parasite reads.

Despite drastic changes in genome architecture in the lineage leading to *Kudoa* and considerable sequence divergence between the two genomes, gene order is highly conserved. Although parasitic cnidarians show rapid protein evolution compared to their free-living relatives, there is limited evidence of less efficient selection. While deleterious substitutions may become fixed at a higher rate, large evolutionary distances between species make robustly analysing patterns of molecular evolution challenging. These observations highlight the importance of filling in taxonomic gaps, to allow a comprehensive assessment of the impacts of parasitism on genome evolution.

## Introduction

Myxozoans are microscopic obligate endoparasites that have a two-stage life cycle, alternating between annelid and vertebrate hosts. Though they were originally classified as protists, molecular data confirm their placement within *Cnidaria* [1, 2]. The extreme morphological simplification observed in the myxozoan lineage is accompanied by drastic shifts in genome composition and structure. Myxozoan genomes are small compared to other Cnidaria, with shortened genes, and lack a large fraction of proteins found in their free-living relatives. Key components of developmental pathways usually required for multicellular development are also missing. In some species, even seemingly essential features such as mitochondria or cytosine methylation have been lost [2–7].

Myxozoans therefore represent an interesting system to examine how shifts in lifestyle impact genome evolution. One interpretation of the changes in genome content is that they reflect adaptations to parasitism [7]. In addition to the changes in genome and gene architecture, parasitic cnidarians also show high rates of protein evolution, tempting speculation about diversifying selection [8–10]. However, not all parasites have compact genomes, and there are multiple routes to a small, fast-evolving genome [4, 11–17]. The potential role of non-adaptive processes, including mutation and drift, should therefore not be discounted. Due to dispersion between hosts, myxozoans may experience recurring population bottlenecks, which could allow deleterious changes to fix more easily [18]. We might therefore ask to what extent the observed patterns are consistent with adaptive streamlining or ineffective selection.

To date, the mechanisms driving sequence evolution in parasitic cnidarians have been difficult to address, given the paucity of suitable data. Limited taxonomic coverage and large evolutionary divergences make obtaining reliable estimates of the strength of selection challenging. In addition, little is known about genome structure, beyond gene content. While chromosomal assemblies are increasingly becoming available for free-living cnidarians [19–25], genomes of comparable contiguity are currently missing for parasitic lineages. Sequencing of organisms like Myxozoa has traditionally been challenging given the minute size of the parasites and their being embedded in host tissue. Many studies only provide transcriptomic data [5, 26–28].

Could recent efforts to sequence metazoan genomes at scale [29] indirectly address this gap? Samples from target species often contain additional sequences from other organisms [30–34]. With the increasing use of long-read sequencing, “contamination” from mutualists and parasites has become easier to detect. The target lists of multiple large-scale genome sequencing projects include likely myxozoan hosts, such as fish and annelids [35–37]. Indeed, several fish samples have shown evidence of infection. Initial draft assemblies of the yellowfin tuna (*Thunnus albacares*) and Atlantic horse mackerel (*Trachurus trachurus*) [38] were found to contain sequences likely derived from myxozoans [31, 32], which were removed from the curated host genome prior to release.

This work examines how we can reconstruct genomes for the myxozoan parasites from sequencing data from infected hosts. In the absence of sufficiently closely related reference sequences, reliably ascertaining which sequences derive from the parasite is challenging. However, we can take advantage of two strategies based on unsupervised learning to address these problems. DNA molecules from the same genome are expected to interact physically within the nucleus (or cell, in the case of prokaryotes). Identifying distinct neighbourhoods in chromatin connectivity networks therefore allows scaffolds from host and parasite to be separated. In addition, sequences from different genomes commonly differ in base and k-mer composition, which allows them to be grouped without explicit knowledge of taxonomy [39–41]. Most genome composition clustering tools are designed for assembled contigs. However, a recently proposed method to generate two-dimensional sequence embeddings with a Variational Autoencoder (VAE) makes clustering of unassembled long reads feasible [32]. This permits strategic assembly of a subset of reads chosen to contain a large fraction of parasite sequences.

Applying these approaches allowed us to generate highquality assemblies for the myxozoans in both *Th. albacares* and *Tr. trachurus*. Phylogenetic analyses identified both parasites as members of the genus *Kudoa*. The assemblies permit fine-grained analysis of genome structure and gene order evolution. Despite the drastic shifts in genome architecture in the lineage leading to *Myxozoa*, the two genomes are highly colinear. Obtaining reliable estimates of the strength of natural selection across large evolutionary distances, including across *Myxozoa*, remains challenging. Past reports of positive selection on specific proteins should therefore be treated with caution. However, tripling the number of available genomes in the family *Kudoidae* makes examining substitution rates on a smaller scale feasible. We observed weak evidence of accelerated protein evolution in *Kudoa* compared to free-living cnidarians, in principle consistent with less efficient selection in parasitic genomes.

## Methods

### Data

Genomic sequences for *Th. albacares* were obtained from muscle tissue from a specimen sampled from the Coral Sea, Australia (fThuAlb1, BioSample SAMEA8654761). The curated *Th. albacares* assembly, generated from PacBio HiFi reads, was previously released under accession GCA_914725855.1. Data for *Tr. trachurus* came from muscle from a specimen sampled from Southampton Water, United Kingdom [38] (fTraTra1, BioSample SAMEA7524396). The *Tr. trachurus* assembly was generated from PacBio single-pass CLR reads, and was previously released under accession GCA_905171665.2. Arima Hi-C data were available for both species.

Additional read sets generated to attempt to increase coverage of the *Kudoa* genomes have been submitted to ENA under run accessions ERR13510304, ERR13510305, and ERR13510306.

### Extracting Kudoa sequences from fish assemblies

#### Identifying sample contamination

Routine curation checks of the initial *Th. albacares* and *Tr. trachurus* assemblies included visual inspection of the scaffolds with BlobToolKit [33]. The observation that a group of low-coverage, GC-poor scaffolds were not assignable to a fish reference genome prompted more detailed assessment. This included examining the taxonomic assignments of all nuclear small subunit ribosomal RNA gene (nSSU) sequences found in the assembly using MarkerScan [31]. Nontarget sequences were removed from the fish assemblies.

As an additional means of visualising potential contamination in the fish samples, two-dimensional representations of sequence tetranucleotide composition were learned by a Variational Autoencoder [32, 42]. This provides a summary of composition that is not solely based on GC content, which may not always be sufficient to distinguish sequences from different sources [43]. The VAE framework can be applied to scaffolds, contigs or long reads (as illustrated in Figure 1 and Figure 4). In addition to allowing visual exploration of both assembled and unassembled sequences, it allows sequences of interest to be extracted. See [32] for a more detailed description. The software required to train the VAE and display read k-mer coverage is available from: https://github.com/CobiontID/read_VAE.

**Fig. 1.**
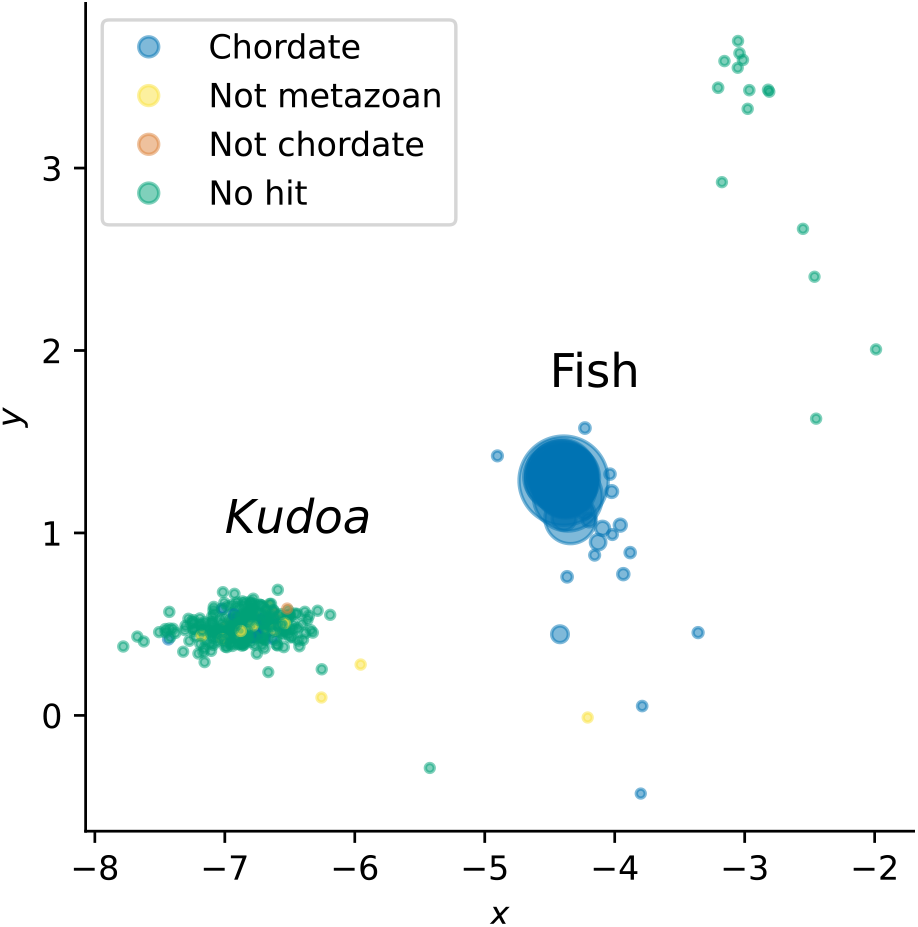
Scaffolds from the preliminary *Th. albacares* assembly prior to decontamination, separated by tetranucleotide composition using a Variational Autoencoder. Each point corresponds to one scaffold, with each axis representing one of two latent dimensions. Size is proportional to scaffold length. The colour of each point indicates the taxonomic assignment of each sequence according to BlobToolKit (*bestsumorder phylum*). In addition to a heterogeneous group of fish sequences on the right, which can all be assigned as chordate, there is a more homogeneous cluster of small scaffolds with unclear taxonomic assignments on the left.

#### Hi-C connectivity

To assess connectivity between scaffolds without manually inspecting Hi-C maps, a simplified representation of the chromatin contact information generated as part of the scaffolding process of the original assemblies was used. For each pair of scaffolds *i* and *j* in an assembly, the number of contacts is encoded in a symmetric *n* × *n* matrix ***A***, where *n* is the number of scaffolds. The matrix entries *a*_*ij*_ and *a*_*ji*_ are incremented by one for each observed connection between *i* and *j*.

The adjacency matrix was next extracted from the connectivity matrix ***A*** and converted into an undirected graph with NetworkX [44]. An adapted version of node2vec was then used to generate node embeddings for each scaffold [45]. The node2vec algorithm performs a biased random walk through the network, and assigns nodes that have similar connectivity patterns similar feature vectors (d = 16), allowing them to be clustered and labelled with KMeans. Since we are interested in detecting groups of scaffolds that form distinct clusters (homophily), biasing the random walk towards visiting nodes it has not traversed before is most appropriate. The parameters *p* and *q* were therefore set to 1 and 0.5, respectively. The code for generating and visualising network embeddings is available from https://github.com/CobiontID/HiC_network.

#### Phylogenetic placement of parasite sequences

Screening of the initial fish assemblies with MarkerScan [31] indicated the presence of nSSU sequences belonging to the family *Kudoidae*. To allow an explicit phylogenetic placement, the nSSU was aligned with nSSU sequences from the SILVA database with SSU-align [46–48]. After masking the alignment with ssu-mask, a tree was inferred using the 16-State secondary structure model in RAxML 8.2.12 [49].

Both trees placed the contaminant from the *Th. albacares* sample with *Kudoa neothunni* and *Kudoa hexapunctata*. To resolve which of the closely related morphotypes the species belongs to, we additionally examined the less conserved nuclear large ribosomal subunit (nLSU) sequence (see [50]), extracted from the original *Kudoa* scaffolds using the eukaryotic nLSU profile (Rfam RF02543). *Kudoa* nLSU sequences from the SILVA database were trimmed with cmsearch [47] to ensure homology, and aligned to the extracted sequences with cmalign. For species outside the clade of interest (species from genus *Kudoa*: *K. iwatai, K. septempunctata, K. unicapsula, K. islandica, K. quadricornis, K. paraquadricornis*), only the longest aligned sequence was retained. Model selection with PHASE 3.0 [51] indicated that the best-fitting model was GTR+G/RNA16A+G, which accounts for secondary structure and dinucleotide substitutions (available in RAxML 8.2.12, [49]). The resulting phylogeny groups the *Kudoa* nLSU found in *Th. albacares* with *K. neothunni* (94% bootstrap support for the bifurcation between *K. hexapunctata* and *K. neothunni*). A simple GTR+G model returned similar results and supports the same conclusion. The placement of the parasite from *Tr. trachurus* is less clear, as there was no sufficiently closely related reference available. This was also the case for the mitochondrial sequence.

#### *Kudoa* genome re-assembly

To produce improved *Kudoa* assemblies, additional HiFi reads were generated from muscle tissue from the same individual fish specimens following standard protocols. However, contamination in these sequencing runs was minimal compared to earlier samples, indicating that the level of contamination is heterogeneous through the fish tissue. Coverage for the *Kudoa* sequences in the newly generated *Tr. trachurus* HiFi data was around 1x, which was insufficient for assembly.

Potential *Kudoa* reads were identified by mapping reads to myxozoan scaffolds from the initial fish assemblies with minimap2 [52], generating VAE embeddings [32], and inspecting the area of the latent space where reads with hits were concentrated. Annotating the embeddings with k-mer coverage additionally allowed discrepancies in coverage between the host and parasite to be visualised.

#### Reassembly from Th. albacares HiFi reads

To account for the large discrepancy in coverage between the fish and the parasite, a new assembly using all available HiFi reads was generated using hifiasm-meta [53]. The resulting contigs were scaffolded with Hi-C data using YaHs [54], which returned a less fragmented assembly than SALSA2 for the same set of contigs [55]. Fish scaffolds were subsequently removed based on tetranucleotide composition.

This captured all scaffolds that belonged to the *Kudoa* connectivity network (node2vec parameters: p = 1, q = 0.5, k = 3, d = 32), as well as some additional short scaffolds with no detectable connections. Mapping the retained scaffolds against the *Th. albacares* genome with minimap2 [52] provided further confirmation that the fish sequences were fully removed.

Visual inspection of Oxford plots of the selected *Kudoa* scaffolds suggested that there was remaining haplotypic variation in the assembly. After purging duplication with purge_dups v. 1.2.5 [56], and manual curation, 40.2 Mb of sequence remained. In addition, a circular contig containing the mitochondrial genome was present.

#### Reassembly from Tr. trachurus CLR reads

Reassembling Falcon-corrected *Tr. trachurus* reads with hifiasm-meta was unsuccessful, presumably due to the relatively high remaining error rate. Reads were therefore instead selected manually based on VAE embeddings, as illustrated in Figure 4. Differences in GC content between the host and parasite contribute substantially to separating the sequences, given the strong correlation between the first latent dimension and GC (*ρ* = 0.9270, *p <* 0.0001, see [32]). In addition, the relatively homogeneous composition of the parasite genome makes it easier to extract reads based on their embeddings. The code used to generate the embeddings and extract reads of interest is available from https://github.com/CobiontID/read_VAE (see [32]).

Wtdbg2 was then used to assemble the pre-selected reads with preset 1, which is designed for reads with high error rates [57]. Mapping the reads back to the assembly suggested that the vast majority of reads in the selected cluster were incorporated into the assembly. Although the majority of host reads were discarded (see Figure 4B), the selection included some fish sequences. Fish contigs were therefore removed as described above for *Th. albacares*. Scaffolding the myxozoan contigs was unsuccessful in this case, likely due to sparse Hi-C signal. After purging with purge_dups the assembly span was close to the haploid size predicted from the k-mer histogram with GeneScopeFK [58, 59]. Low coverage made fitting the model reliably challenging, resulting in some uncertainty. A more stringent alignment score threshold (-a 90) was necessary to avoid over-purging. Consensus quality values for both assemblies were calculated with Merqury.FK [60].

#### Assessing genome structure

After preliminary inspection with PretextMap [61], HoloViews and Datashader were used to visualise chromatin connection data extracted from a Cooler file [62]. Code for viewing the map is available from https://github.com/CobiontID/HiC_network. Putative telomeric repeats were identified by screening the assemblies with https://github.com/tolkit/telomeric-identifier.

### Gene annotation

#### Nuclear gene annotations

BUSCO v5.3.2 [63] with metazoa_odb10 was used to allow an initial if limited assessment of the gene content of the assemblies. To obtain a more comprehensive annotation, BRAKER3 [64] was run with protein hints from *K. iwatai* and uniprot proteins for *Cnidaria* (downloaded 25/08/2023). Although RNA-seq data were available for *Th. albacares*, the fraction of reads aligning to *Kudoa* was insufficient to allow transcript-based annotation. Both genomes were masked using RepeatModeler and RepeatMasker (docker://dfam/tetools:latest, downloaded 28/09/2023) before being annotated. The AUGUSTUS configurations for each species that were generated as part of the BRAKER3 pipeline were subsequently used to improve retrieval of BUSCO genes. Annotations are available from https://github.com/CobiontID/Kudoa_genomes.

#### Mitochondrial annotation

The meta-assembly from the topped up *Th. albacares* read set contained an 18, 219 bp contig with 92.23% nucleotide similarity to the *K. hexapunctata* mitochondrion, which is of similar length. There was no full *K. neothunni* mitochondrial sequence available at the time of writing. For *K. sp. trachurus*, retrieving HiFi reads that aligned to Cox1 in *K. hexapunctata* (YP_009158818.1) with miniprot [65] and assembling them with hifiasm-meta yielded a linear 22, 958 bp contig. Mitochondrial gene annotations were obtained with MITOS2 [66], using translation table 4.

### Comparative analyses

Several new annotated chromosomal assemblies from free-living cnidarians provided an opportunity to examine sequence evolution across the phylum, namely *Blastomussa wellsi, Galaxea fascicularis, Montipora capitata, Nematostella vectensis, Porites lutea*, and *Ricordea florida* (annotations available from https://github.com/Aquatic-Symbiosis-Genomics-Project/cnidaria_annotations). In addition, the dataset included previously published sequences from *Acropora millepora, Actinia tenebrosa, Ceratonova shasta, Exaiptasia diaphana, Henneguya salminicola, Hydra vulgaris, Hydractinia symbiolongicarpus, Kudoa iwatai, Myxobolus honghuensis, Myxobolus squamalis, Orbicella faveolata, Pocillopora damicornis, Polypodium hydriforme, Sphaerospora molnari, Stylopora pistillata, Tetracapsuloides bryosalmonae*, and *Thelohanellus kitauei*. Free-living species with chromosomal assemblies were favoured in compiling the dataset. Sources for publicly available data are shown in Table S1.

For parasitic cnidarian species where transcripts but not coding sequences were available (*C. shasta, K. iwatai, P. hydriforme, S. molnari, T. bryosalmonae*), TransDecoder [67] was used to identify the longest predicted coding sequences, which were then filtered with CD-HIT [68] to remove identical sequences.

OrthoFinder2 [69] was used to identify orthogroups with default settings. The annelid *Lumbricus rubellus* was included as an outgroup [70]. Amino acid sequences for orthogroups containing all species were aligned with MAFFT v. 7.481 [71]. Subtrees of 1:1 orthologs found across all species were extracted with PhyloPyPruner [72], giving a total of 95 alignments (settings –mask longest –min-pdist 2e-8). A second, less conserved set of orthologs containing at minimum all corals and all three *Kudoa* species was also compiled (n = 491).

### Substitution rates

The site heterogeneous PMSF model in IQ-TREE2 was used to infer a supermatrix tree [73, 74]. Given the long evolutionary distances between some of the included species, site heterogeneous models are expected to be more robust against long branch artefacts [75]. The topology returned the expected species relationships.

To estimate codon substitution rates, orthologous coding sequences were aligned with PRANK in “codon” mode [76]. The algorithm considers phylogeny in order to distinguish between insertions and deletions. This should, in principle, allow more reliable alignments given the extensive loss of sequence in the myxozoan lineage (parasitic sequences contained a larger fraction of alignment gaps). In addition, PRANK provides information about inferred insertion and deletion events for each branch. The PMSF species tree was used as a starting tree for alignment. Several of the coding sequences obtained from public databases contained internal frameshifts. To maximise the number of usable sequence alignments, the affected alignments were therefore processed with OMM_MACSE [77], using PRANK and the abovementioned settings. All other filtering steps were switched off.

To test whether there is evidence of less effective selection on the myxozoan branch, codeml was run for each alignment [78] with M0, which infers a single value of *ω* for the whole phylogeny, as well as a clade model (M2), where separate estimates of *ω* are obtained for different pre-defined groups of branches.

Inspection of initial parameter estimates indicated that comparisons between the myxozoan clade and free-living species are potentially unreliable. Estimates of the rate of synonymous divergence, *dS*, commonly exceed 3 for multiple branches (Figure S13), which suggests potential issues with saturation [79]. Since breaking up long internal branches by adding more myxozoan species to the alignment was not possible, we can instead consider the *Kudoa* clade alone. Since the group including corals and sea anemones had more manageable branch lengths, they were chosen to represent free-living species. Estimating *ω* for the non-myxozoan parasite *P. hydriforme* was not possible, given the branch length. Missing data were treated as ambiguous, and alignment columns where at least one *Kudoa* species had a gap were removed.

In addition to *ω*, the ratios of conservative or radical amino acid substitutions over synonymous substitutions, *ω*_*C*_ and *ω*_*R*_, were estimated for the clades of interest. Here, substitutions affecting the polarity or volume of the residue are considered “radical” [80]. The procedure for fitting the “Conservative or Radical” model with bppml [81] is described in https://github.com/claudia-c-weber/CoRa.

## Results

### Identifying myxozoan contamination in two fish

Curation checks of the initial *Th. albacares* assembly highlighted a number of scaffolds with no clear taxonomic assignment, which were compositionally distinct from those assigned to the target genome (Figure 1). Multiple scaffolds in the cluster contained nSSU and nLSU loci that suggested a myxozoan origin. Phylogenetic placement identified *Kudoa neothunni*, which is known to infect *Th. albacares*, as the likely source of the sequences [50, 82]. Given the aim of identifying additional *Kudoa* sequences, we next examined if the remaining scaffolds in the cluster belong to the same genome.

Recording chromatin contacts between each pair of scaffolds and examining the resulting connectivity graph allows sequences from different genomes to be distinguished. Groups of highly interconnected (homophilous) scaffolds can be identified and labelled by performing a biased random walk over the graph with node2vec [45]. For the *Th. albacares* data, the network shows clear separation between the fish genome and a cluster of scaffolds containing *Kudoa* nSSU and nLSU sequences (Figure 2).

**Fig. 2.**
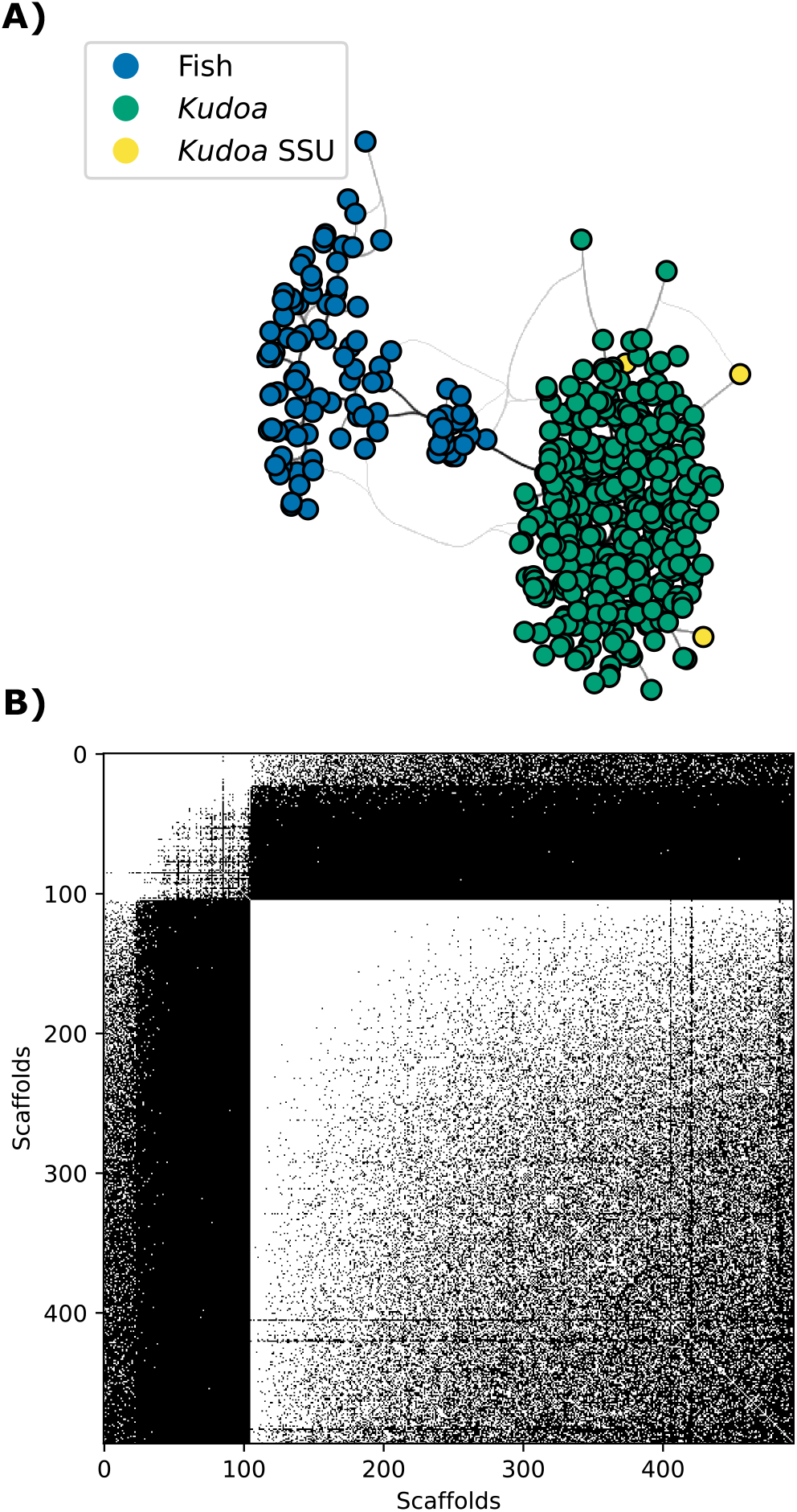
Chromatin contact data from the initial assembly of *Th. albacares* reveals the presence of two genomes. **A)** The connectivity network displays distinct components. Each point represents one scaffold, and the lines show connections between scaffolds shaded according to weight. The scaffolds in the group on the right that are marked in yellow contain the *Kudoa* nSSU locus. The remaining colours represent cluster labels from node2vec network embeddings [45] **B)** A binary adjacency matrix shows connections between pairs of scaffolds. White pixels represent the presence of at least one connection and black pixels represent the absence of connections. Scaffolds are sorted by cluster membership, revealing two distinct groups (*Kudoa* scaffolds appear in the bottom right corner). A subset of fish scaffolds show spurious connections with *Kudoa* scaffolds.

The *Kudoa* network cluster overlaps almost perfectly with the group of compositionally distinct scaffolds in Figure 1, confirming that these sequences are derived from the same source. The sole exception is a scaffold containing *Kudoa* mitochondrial sequence, which shows evidence of interaction with the *Kudoa* nuclear genome, but does not otherwise resemble it. Crucially, the labels assigned by network analysis also correspond to manual assignments made by an expert curator, and were used to remove additional small *Kudoa* scaffolds from the host assembly. Note that although a *K. iwatai* genome is available the corresponding proteins are not present in the NCBI database. An unsupervised approach was therefore necessary for the initial assessment.

A similar pattern was observed in the initial assembly of *Tr. trachurus* (Figure S1, Figure S5). Here, the phylogenetic placement of the ribosomal marker sequences was less clear, and previous reports suggest that the fish can be infected with multiple species of *Kudoa* (see Figure S2, S3). The parasite detected in *Tr. trachurus* is hereafter referred to as *Kudoa sp. trachurus*. Having confirmed that the contamination in both fish assemblies resulted from infection by *Kudoa*, we next asked whether the extracted scaffolds contain complete genomes.

The span of the *K. neothunni* scaffolds was only slightly below that of the published assembly of *K. iwatai* [2], the most closely related available genome. However, the span of the scaffolds assigned to *K. sp. trachurus* was substantially smaller than either of the others. Retention of widely conserved genes was also lower (Table 1, Figure 3). In addition, a comparison of *K. neothunni* and *K. sp. trachurus* suggested that multiple stretches of homologous sequence were missing in the latter. Both new initial assemblies nevertheless showed higher contiguity. We therefore returned to the unassembled read data, to recover additional parasite reads and produce improved genome assemblies.

**Table 1.** Statistics describing *Kudoa* genome assemblies after purging duplication and removing fish sequences. Details for *K. neothunni* refer to the curated assembly. *K. iwatai*, two other myxozoans with scaffold N50 > 5 kb, and the free-living cnidarian *N. vectensis* are included for comparison. BUSCO completeness is based on metazoa_odb10 (*n* = 954). Completeness estimates for public myxozoan assemblies may be reduced due to low contiguity. *Contig N50. **Statistics from NCBI.

| Species | Assembly | Assembly span | Scaffold N50 | BUSCO complete |
| --- | --- | --- | --- | --- |
| *K. neothunni* | Initial | 27.4 Mb | 106.8 kb | 29.1% |
|  | Final | 40.2 Mb | 12.0 Mb | 40.0% |
| *K. sp. trachurus* | Initial | 19.1 Mb | 472.0 kb | 24.0% |
|  | Final | 35.4 Mb | 2.8 Mb* | 39.7% |
| *K. iwatai* ** | GCA_001407235.2 | 31.2 Mb | 40.2 kb | 28.5% |
| *H. salminicola* ** | GCA_009887335.1 | 61.4 Mb | 7.6 kb | 26.8% |
| *T. kitauei* ** | GCA_000827895.1 | 150.5 Mb | 149.8 kb | 25.1% |
| *N. vectensis* ** | GCA_932526225.1 | 269.4 Mb | 17.9 Mb | 96.9% |

**Fig. 3.**
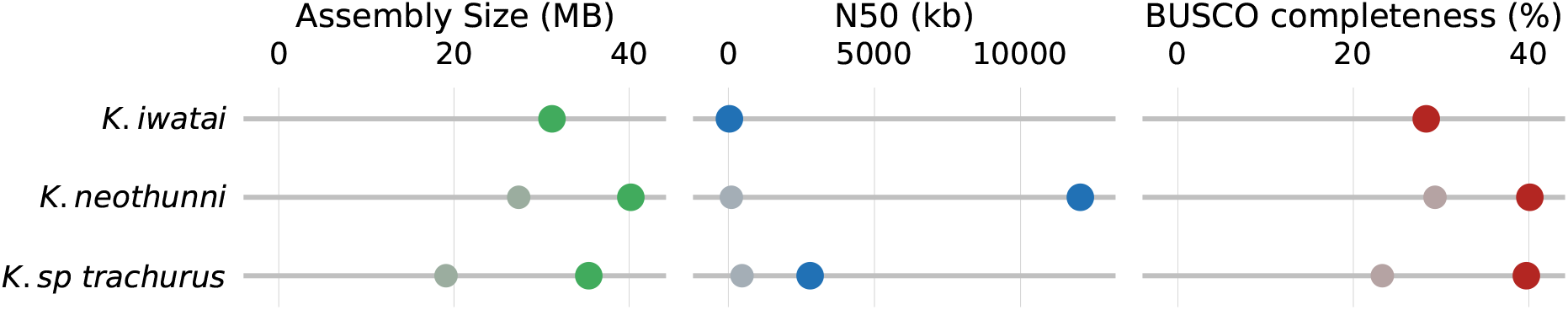
*Kudoa* assembly statistics compared to *K. iwatai*, before and after reassembly. Smaller greyed out points represent the originally retrieved scaffolds. Coloured points represent reassembled, purged genomes. For the *K. neothunni* reassembly statistics reflect the curated genome. BUSCO completeness is based on metazoa_odb10.

### A near-chromosomal assembly despite low-coverage

As shown in Figure 1, differences in composition allow sequences from the host and the parasite to be distinguished [32]. Examining two-dimensional read embeddings can therefore provide an overview of the relative abundance of potential *Kudoa* reads, including those not yet incorporated into an assembly (Figure 4).

**Fig. 4.**
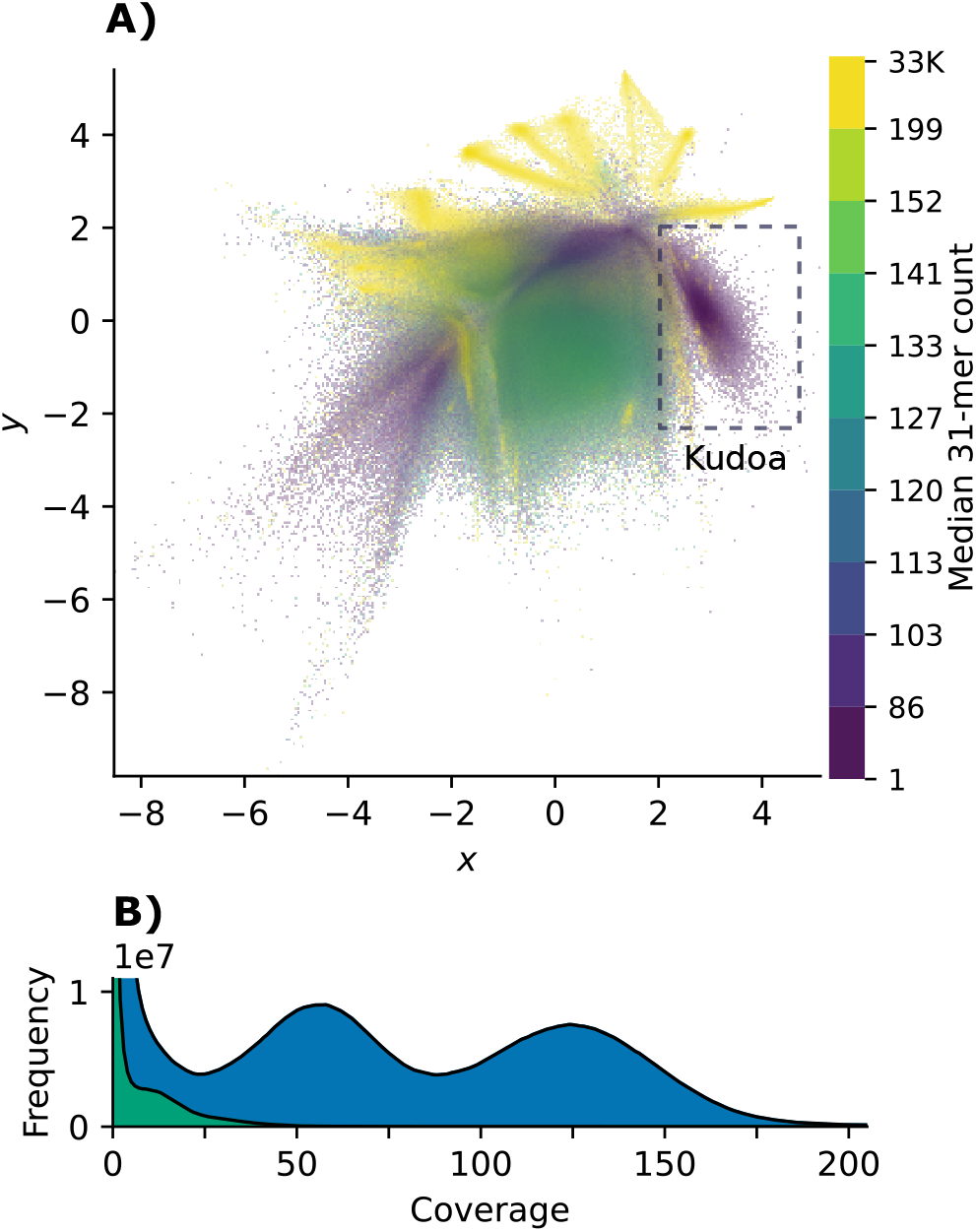
**A)** shows 2D VAE embeddings of reads from *Tr. trachurus*, coloured by k-mer coverage (k = 31). The cluster containing reads that map to *K. sp. trachurus* scaffolds, highlighted on the right, has markedly lower coverage than the host. The rectangle indicates the region of the latent space containing the reads selected for targeted assembly. **B)** shows the k-mer count distribution (k = 31) of the full *Tr. trachurus* read set (upper curve, shaded blue), with the distribution of the reads in the selection superimposed (lower curve, shaded green). The latter highlights the coverage peak of the *Kudoa* sequences.

The k-mer coverage for reads mapping to the initial *K. neothunni* scaffolds was around seven (see Figure S6). This is below the recommended sequencing depth for *de novo* genome assembly from HiFi data. We therefore generated additional long read data from the same *Th. albacares* individual. However, the majority of putative *Kudoa* reads in the combined dataset came from the first extraction (22,263 of 27,248), and only 390 were added by the top-up run. In the case of *K. sp. trachurus*, which was initially assembled from CLR reads from *Tr. trachurus*, haploid k-mer coverage was around 14 (Figure 4). Additional HiFi reads from the same fish again yielded few *Kudoa* reads. The relative paucity of myxozoan sequences noted in both cases is consistent with low-level infection that was not conspicuous during sample processing.

Despite limited coverage, it was straightforward to produce a highly contiguous genome for *K. neothunni* from a standard meta-assembly of all HiFi reads from *Th. albacares*, followed by scaffolding (see Methods). Assembly size, contiguity, and completeness all improved compared to both the initial assembly and *K. iwatai* (Table 1, Figure 3).

Meta-assembly was not feasible for the CLR data available for *Tr. trachurus*. An alternative approach to addressing discrepancies in coverage was therefore necessary. Here, removing the majority of fish reads by tetranucleotide composition with the help of VAE embeddings permitted a more contiguous and complete assembly of *K. sp. trachurus* (see Figure 4; Table 1). While it is possible that relying on composition alone to pre-select reads could lead to some parasite sequence being missed, very few reads outside the selected region of the embedding space credibly mapped to the previously extracted *K. sp. trachurus* scaffolds (n = 592).

Though the assembly quality metrics for *K. sp. trachurus* do not match those of *K. neothunni*, the genome size is consistent with predictions from the k-mer distribution. Given the limited coverage for *K. neothunni*, it was not possible to reliably estimate the expected genome size. The relatively lower quality score for *K. sp. trachurus* is in line with the more fragmented assembly. The larger spans of both assemblies compared to *K. iwatai* are perhaps due to genomic regions that are difficult to assemble from short reads being absent in the latter.

The remarkable contiguity of the *K. neothunni* assembly provides insight into its genome structure. After manual curation, three large scaffolds representing chromosomes were obtained (Figure 5). One chromosome end contained tandem repeats indicating the presence of a telomere (see Supplementary Material). The observed motif, (TAAACC)_*n*_, differs from the canonical metazoan motif seen in free-living *Cnidaria* by a single nucleotide [83, 84]. Telomerase reverse transcriptase was also retrieved from the genome. Although the telomeric motif was observed in the reads used to assemble *K. sp. trachurus*, it was not present in the assembly, presumably reflecting limitations of the sequencing technology used. The observed haplotypic duplication in both species is consistent with the expectation that the fish-infecting stage of the parasite is diploid [8]. A circular mitochondrial genome was also present in *K. neothunni*, consistent with previous reports from myxozoans [85]. Analysis of the Cox1 gene provided further confirmation that the *Th. albacares* specimen was infected with *K. neothunni* (Figure S4).

**Fig. 5.**
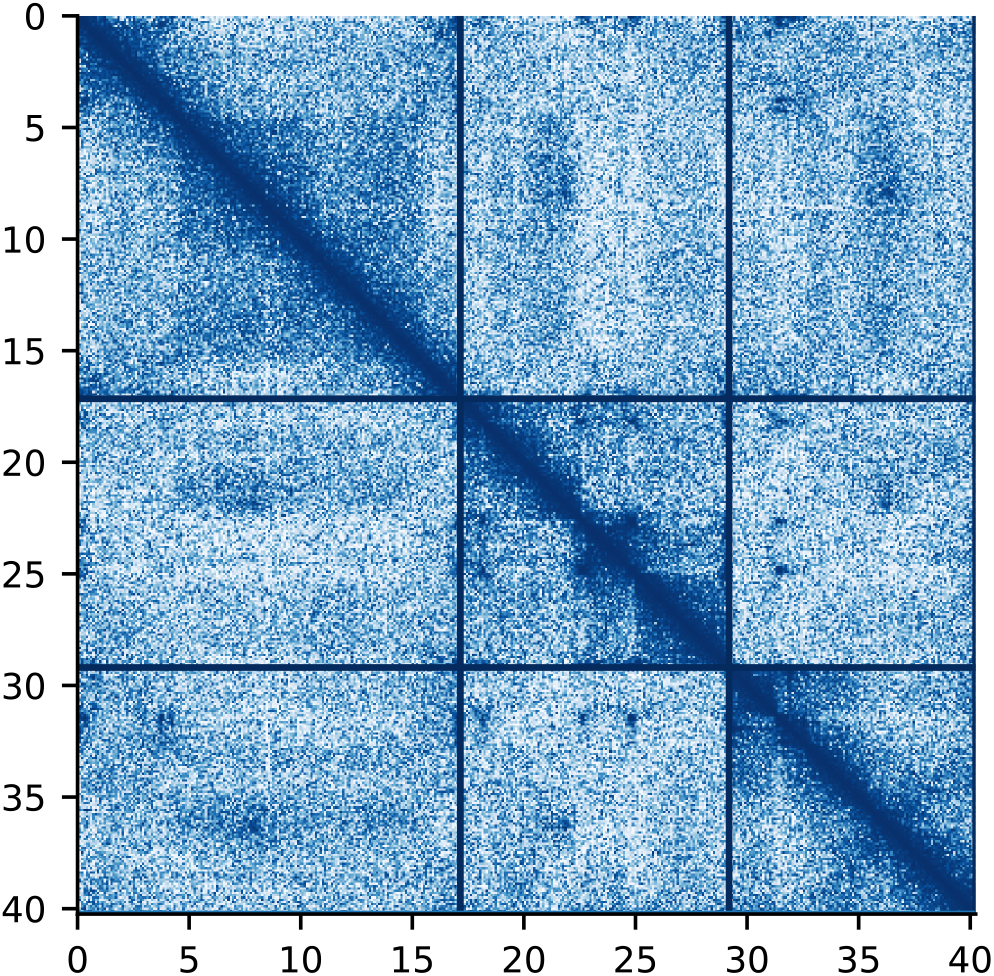
Hi-C map for the curated *K. neothunni* nuclear genome assembly. Darker shading indicates stronger interactions. Chromosomes are ordered by size and demarcated by grid lines.

### Reduced, but not streamlined

Gene annotation confirms that the two new *Kudoa* genomes have a very reduced repertoire of protein coding genes (Table 2), in line with previous reports from *K. iwatai* [2]. As very few of the transcriptomic reads generated for *Th. albacares* mapped to *K. neothunni*, gene predictions are based on protein hints from *K. iwatai* and other cnidarians (see Methods).

**Table 2.**
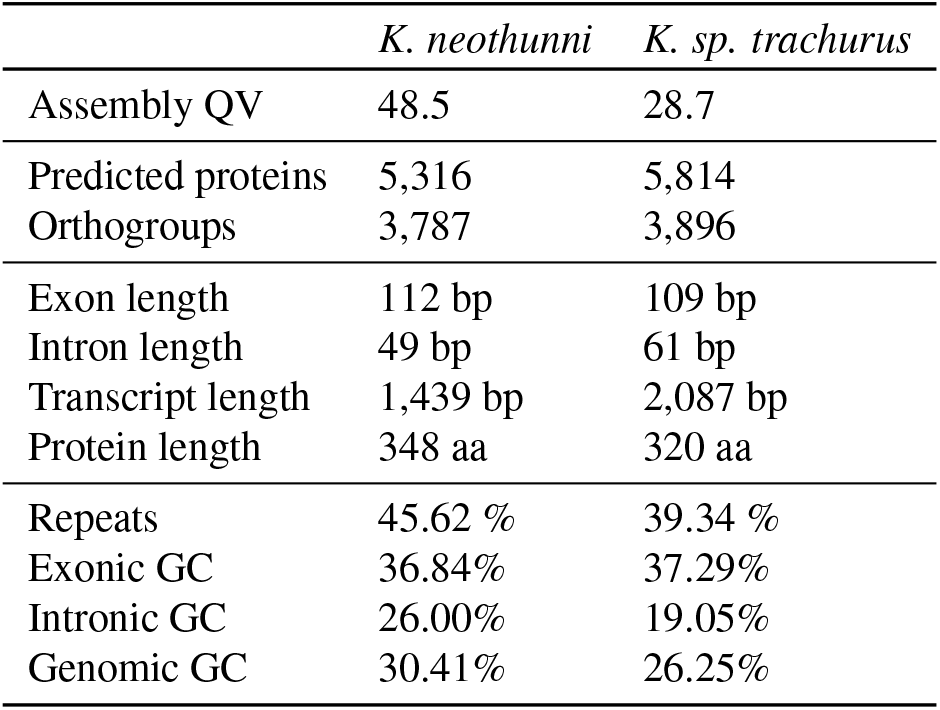
Annotation statistics for purged assemblies. Genes were predicted with BRAKER3. Feature lengths include alternative transcripts. Transcript lengths refer to unspliced sequence features including exons and introns. Repeat statistics from RepeatMasker. The consensus quality value (QV) was calculated with Merqury.FK for for k = 21.

Divergence between *K. iwatai* and the two more closely related new genomes is substantial, as illustrated by the species tree in Figure 6. Some turnover in gene content is therefore expected, and it is not surprising that a subset of *K. iwatai* proteins have no matches in the new *Kudoa* genomes. Of the transcripts reported for *K. iwatai*, 73% have orthologs in *K. neothunni* and 74% in *K. sp. trachurus*. Meanwhile *K. neothunni* and *K. sp. trachurus* show overlaps of 78-83%, with 66-69% of their predicted genes having orthologs assigned in *K. iwatai*. In general, *K. iwatai* shares a smaller number of orthogroups with the other species in the dataset than the other two *Kudoa* genomes do. This suggests that the new genomes are largely intact. Differences in the annotation process may also account for some of the discrepancy. Annotations based on protein hints might be biased towards producing predictions with detectable sequence homology. On the other hand, myxozoans are prone to intron retention [4], which may lead to difficulty identifying homologs for *K. iwatai* transcripts.

**Fig. 6.**
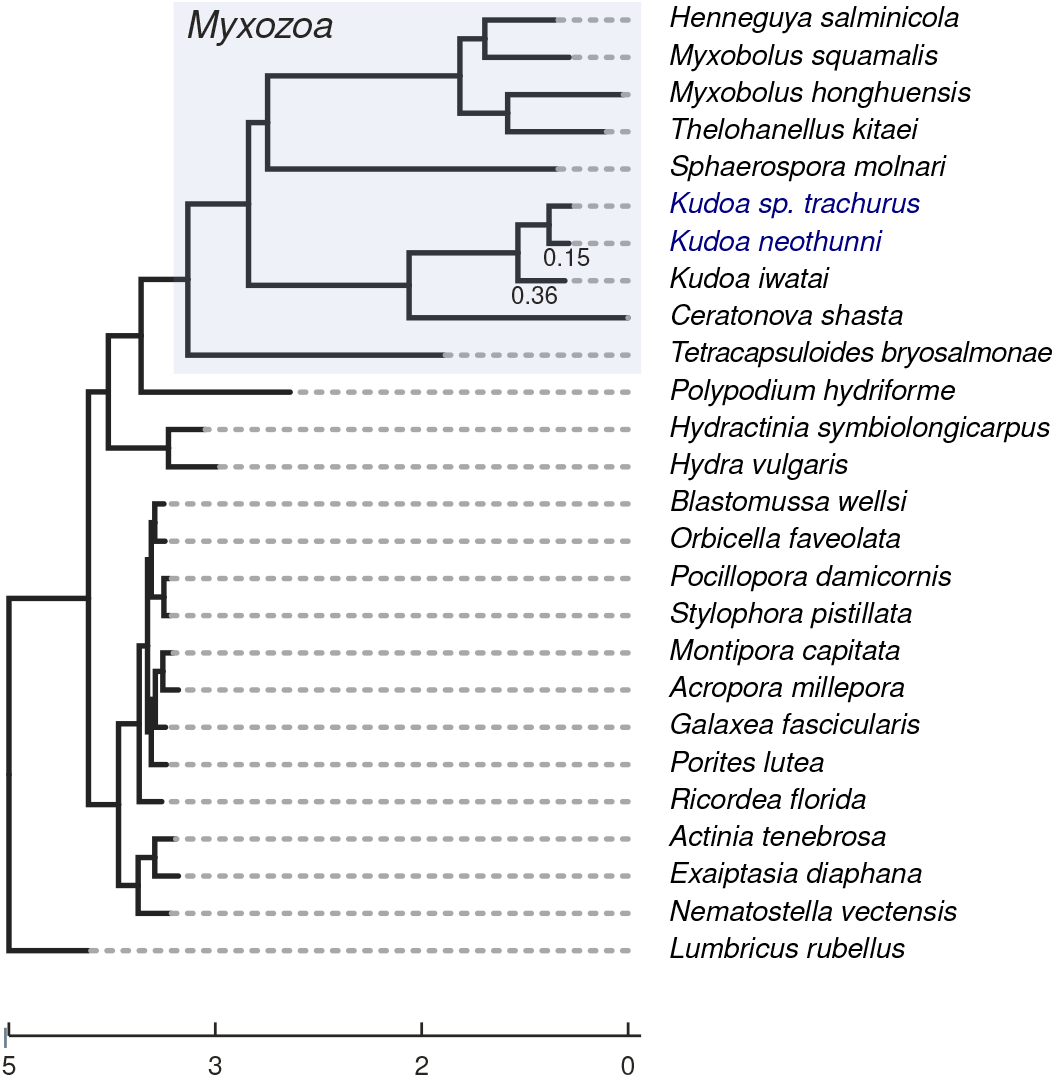
Supermatrix tree illustrating cnidarian species relationships, inferred under the PMSF model using 95 orthologs found in all species, and rooted using the outgroup, *Lumbricus rubellus*. Branch lengths indicate amino acid distances. The myxozoan clade is highlighted in grey. The two new *Kudoa* genomes are shown in blue. Inferred branch lengths for the *Kudoa* clade are shown to give an indication of the level of divergence. The topology is consistent with myxozoans being most closely related to *Medusozoa*. All splits have 100% bootstrap support, except the node at the base of *S. molnari* (80% support).

Other correlates of genome contraction are also broadly conserved across the group (Table 2). *Kudoa* genes are comparatively short, with a median protein length of 320-348 amino acids, compared to 456 amino acids in the free-living species *N. vectensis*. Median exon lengths are consistent with reports from *K. iwatai* [2], and show similar distributions in both species (*K. sp. trachurus* median = 109 bp; *K. neothunni* median = 112 bp; Figure S7). Interestingly, *K. neothunni*, which has the larger assembly, has slightly longer proteins and a lower inferred rate of deletion in orthologous coding sequences than *K. sp. trachurus*. As with other myxozoans, genomic GC content is relatively low (Figure S9). Meanwhile, *K. sp. trachurus* appears to have longer, more AT- and repeat-rich introns than *K. neothunni*, while exonic base composition is similar (Figure S7, S8).

Just under half of the sequence in both genomes is predicted to be repetitive, with *K. neothunni* containing a higher percentage of older repeats, as well as more repeats in general (Table 2, Figure S11). Given the differences in sequencing technology and contiguity, some of this difference may be artefactual. The median distance between annotated genes across all chromosomes in *K. neothunni* is 3,578 bp. *K. sp. trachurus* is more gene dense, with a median intergenic distance of 2,363 bp across contigs, in line with its lower repeat content. Hence, although the *Kudoa* genome is relatively small, it is not as gene dense as those of some other metazoan parasites [86], nor is it depleted in repeats.

### Extensive gene order conservation in *Kudoa*

The contiguity of the new *Kudoa* genomes allows us to examine gene order conservation. With the exception of several inversions, they are remarkably colinear (Figure 7). The overall rank correlation between the adjusted midpoint locations for all orthologs found on the *K. neothunni* chromosomes is *ρ* = 0.8789 (Pearson’s correlation coefficient = 0.8750). This is initially perhaps surprising, given reports from free-living cnidarians, which show conservation of macrosynteny, but not microsynteny [25]. For example, fine-scale gene order conservation is lost between *H. symbiolongicarpus* and *H. vulgaris* [19]. Though these hydrozoan species are more evolutionarily distant from each other than *K. neothunni* and *K. sp. trachurus*, the difference in branch length (*t*) is only around two-fold (*t* = 0.66 versus *t* = 0.32; see Figure 6). In addition, comparisons between free-living species pairs with similar inferred levels of amino acid divergence, such as *B. wellsi* and *M. capitata* (*t* = 0.32), also reveal less fine-scale conservation overall. Note, however, that this pattern varies by chromosome (see Figure S12). There was no evidence that the three inferred chromosomes in *K. neothunni* show any correspondence to linkage groups in *N. vectensis* – as one might expect given the extent of ancestral gene loss in *Myxozoa*. However, these results suggest that there is little indication of ongoing rapid turnover in gene order.

**Fig. 7.**
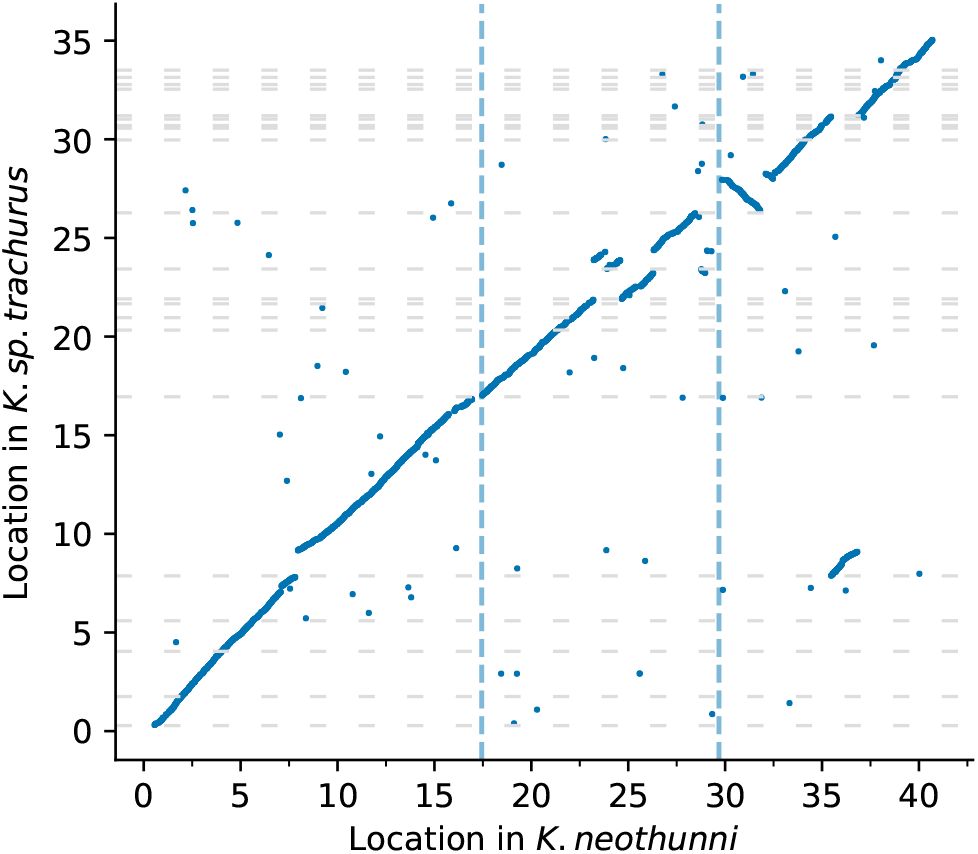
Midpoint locations of 1:1 orthologs, with *K. neothunni* chromosomes ordered from largest to smallest along the x-axis. Dashed vertical lines indicate chromosome boundaries, and dashed horizontal lines show contig boundaries. For the purpose of visualisation, *K. sp. trachurus* contigs are ordered and oriented based on the locations of the corresponding orthologs in *K. neothunni*.

### Is protein evolution accelerated in myxozoans relative to free-living cnidarians?

Given the possibility that myxozoans are subject to transmission bottlenecks [18], we might ask to what extent a propensity for drastic gene loss, gene contraction and shifts in genome architecture correlate with (slightly) deleterious changes reaching fixation. While we cannot conclusively examine gene turnover given the currently available genome annotations, rates of protein evolution may provide an indication of whether natural selection is less effective in the myxozoan lineage. The myxozoan branches are very long, even relative to the distance between the annelid *L. rubellus* and its most recent common ancestor with *Cnidaria* (*t* = 1.23, see Figure 6), in line with previous reports. The observed pattern appears consistent with accelerated protein evolution in the parasitic lineages compared to their freeliving relatives, though it is not clear to what extent this might be influenced by generation time or mutation rate.

The rate ratio of nonsynonymous to synonymous substitutions, *ω* = *dN/dS*, provides a measure of the relative strength and direction of protein-level selection. When most changes are assumed to be deleterious, larger *ω* values can be interpreted as evidence of less effective selection. In this case, we might expect that myxozoan sequences show slightly higher ratios than sequences from non-parasitic species. Given large evolutionary distances and excessive synonymous divergence, it was not feasible to obtain reliable parameter estimates across the whole myxozoan clade (Figure S13). We can therefore consider the most densely sampled part of the myxozoan tree, and compare rates of evolution between *Kudoa* and free-living cnidarians – specifically subphylum Anthozoa (sea anemones and corals), which are well-represented in the dataset. The results suggest that selection is primarily negative (*ω* ≪ 1). Figure 8 shows that *ω* is slightly lower in free-living cnidarians than in *Kudoa* for the majority of protein coding sequences, in principle consistent with modestly weaker negative selection in parasitic species. A similar pattern was apparent when radical and conservative amino acid substitutions were considered separately (see Figure S14).

**Fig. 8.**
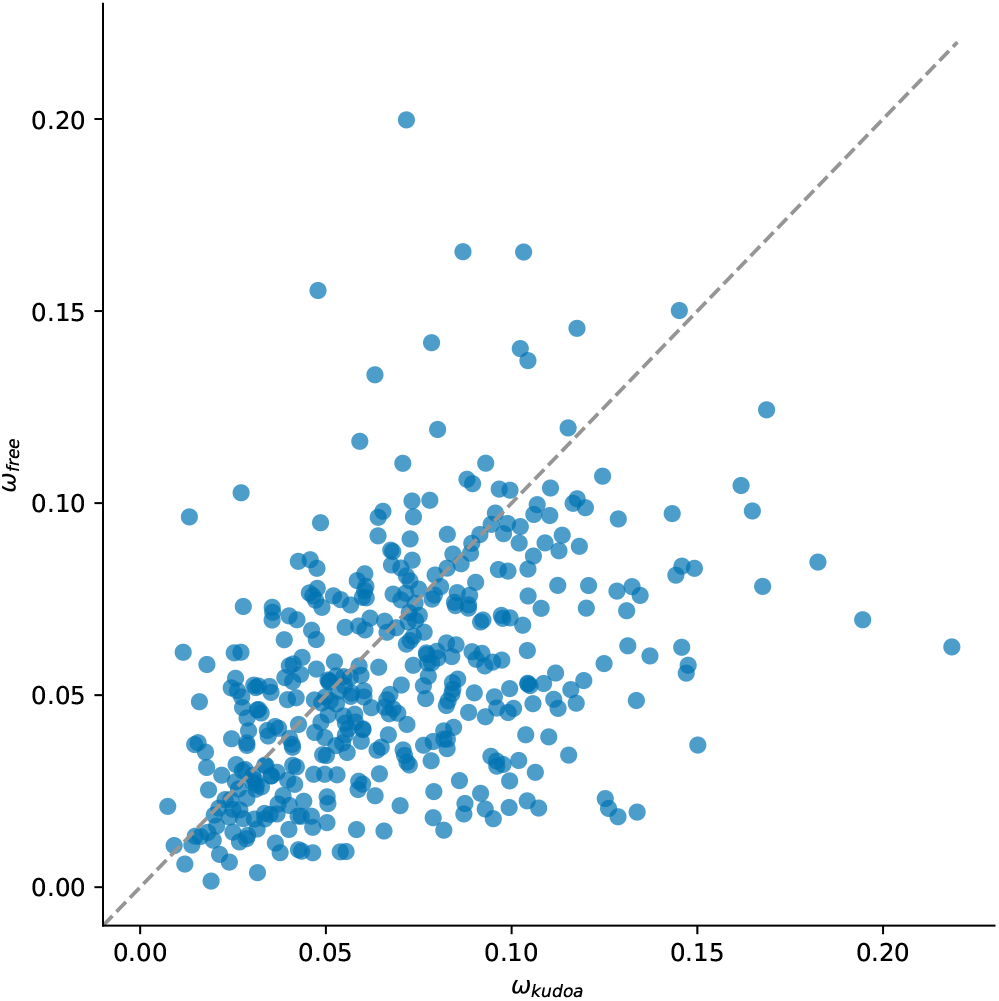
*Kudoa* show weak evidence of accelerated protein evolution compared to free-living cnidarians. Each point represents one alignment (n = 419). The x-axis shows estimates of *ω* for the *Kudoa* lineage, while the y-axis shows estimates for the free-living lineage (Anthozoa) under M2 for the same alignment.

Alignment columns with gaps in any of the three *Kudoa* species were not considered in this comparison. Myxozoan sequences contained a larger proportion of gaps overall, especially for sites present in at least 50% of species. If more dispensable parts of proteins are preferentially lost in parasitic lineages, the corresponding sequences in free-living species are likely subject to less selective constraint on average, potentially masking differences between the groups. In line with this prediction, *ω* across the tree is larger when all columns are retained (median = 0.0686 without filtering, versus 0.0481). When only *Kudoa* sequences are considered, *ω* estimates are broadly similar, suggesting that taxonomically less conserved proteins show comparable levels of negative selection (median = 0.0583, *n* = 2, 588). Note that we do not expect different subsets of genes to be directly comparable in terms of parameter estimates.

These observations suggest that *Kudoa* proteins are relatively constrained, though they may fix amino acid changes somewhat more readily than their free-living relatives. This is not explained by the myxozoan ancestor being excluded from the foreground branch. Despite statistical limitations compromising the accuracy of the estimates, there was no indication that *ω* is large across Myxozoa as a whole.

## Discussion

This work illustrates how eukaryotic parasite genomes can be retrieved from low-level contamination from host samples. Two infected fish each yielded genomes from previously unsequenced *Kudoa* species. Combining information about physical interactions and sequence composition permitted grouping and taxonomic assignment of sequences, despite limited information from conserved markers. The benefits of unsupervised approaches, which can learn the structure of a dataset from unlabelled inputs, are especially apparent in myxozoans. Given high substitution rates [8, 9] and a limited availability of sufficiently closely related annotated reference genomes, reliable labels are scarce. Beyond recovering cobiont genomes from mixtures of sequences, the process also serves to decontaminate the target genome.

The strategy outlined here is not intended to provide a onestep solution to producing complete, separated genomes from samples containing multiple organisms. In many instances, common workflows for assembly and scaffolding that already make use of chromatin contacts will be sufficient to provide a starting point for genome curation [54, 55, 87, 88]. Instead, this work demonstrates how less straight-forward situations can be approached. The removal of additional *Kudoa* sequence from an updated release of the *Th. albacares* genome based on the analyses presented here is a point in case. Though the majority of myxozoan scaffolds were correctly identified from chromatin maps, network embeddings had an advantage in identifying less conspicuous short fragments. Similarly, pre-selecting reads will often be unnecessary, and could lead to sequences from the organism of interest being excluded, but proved useful for the *Tr. trachurus* dataset.

In the two examples shown, both sources of evidence were consistent with the same species assignments, though they offer different advantages. Larger genomes can display considerable compositional heterogeneity between regions (see [32]). The chromatin network embeddings therefore fared better at grouping host scaffolds together than embeddings based on sequence composition. Meanwhile, the *Kudoa* sequences’ overall reduced GC content results in a compact latent representation, making both approaches suitable for retrieving them. This situation is likely to be mirrored for many other host-parasite associations where the parasite has a reduced, compositionally extreme genome [89]. On the other hand, compressed tetranucleotide representations learned by a VAE are useful for extracting sequences of interest from unassembled read sets (though they cannot make use of sequence information as effectively as an assembler). In principle, it would also be possible to consider chromatin interactions and sequence composition simultaneously, for example using latent variable models that operate on graphs [90]. However, initial experiments showed no advantage over node2vec, perhaps because tuning VAEs to produce useful feature embeddings for small sets of contigs is not straightforward [32]. A further option would be to incorporate information from the assembly graph (see [41]), although performance on non-microbial genomes would need to be explored.

What do the new *Kudoa* genomes reveal about myxozoan evolution? Although there are now multiple myxozoans that have been sequenced using long-read technology [91], the near-chromosomal *K. neothunni* genome is, thus far, an outlier. The high contiguity of both *Kudoa* assemblies allows us to more easily examine aspects of genome organisation, such as coding density, the repeat landscape, and patterns of rearrangement. It also provides clues about the karyotype, which has not been directly observed to date. The data are consistent with the presence of three chromosomes in *K. neothunni* - four were reported in the myxobolid *H. gigantea* [92]. Given a pattern of shared breaks (Figure 7), we might speculate that *K. sp. trachurus* also has at least three chromosomes. Despite a limited gene repertoire, *Kudoa* genomes do not appear especially compact compared to some other metazoan parasites [86, 93]. Of course, the proteome remains the most obvious source of information about possible functional changes.

It is tempting to look for evidence of adaptive evolution in the coding sequences of parasitic cnidarians, given drastic changes in lifestyle. Further, diversifying selection driven by host-parasite interactions ought to be relatively easy to detect, given that the pressure to adapt is expected to persist over time [79, 94, 95]. However, it is also apparent that sparse taxon sampling leads to difficulties in inferring the strength of selection across the Myxozoa. Although there have been reports of positive selection in myxozoan genes [91, 96], they are based on comparisons over large evolutionary distances. The problems with estimating evolutionary parameters observed in this work for comparable levels of divergence suggest that these analyses may be prone to statistical artefacts, and should therefore be interpreted with caution. Alignment error, which tends to increase with divergence, is also a concern for detecting positive selection [79, 97, 98].

The prevailing direction of inferred selection across Cnidaria is negative, as is commonly observed for phylogenomic datasets (see [80], Table 1), with signs of a small increase in *ω* in *Kudoa*. This accords with past reports from mitochondrial sequences in *Kudoa* [10] (but see [99]), as well as theoretical expectations under less effective selection [100]. However, the magnitude of the difference in selection efficiency between parasitic and free-living species is challenging to determine. Large discrepancies in branch lengths between different parts of the tree could confound analyses of codon substitution patterns. It has been noted that sequence divergence correlates negatively with *ω* estimates, with long distances being especially problematic [101, 102]. Therefore, it is theoretically possible that the values for *Kudoa* are systematically underestimated relative to the true level of negative selection, obscuring the extent of rate acceleration relative to free-living species. It is difficult to assess how large this effect would be in practice. In addition, myxozoan genomes have, on average, lower GC content, violating the stationary assumption and potentially affecting substitution rates. As a result, our ability to draw firm conclusions about the efficacy of selection in myxozoans remains limited.

The pervasive loss of coding sequence in myxozoans also raises questions about the limitations of conventional estimates of protein-level selection. Measures of the strength of selection such as *ω* do not consider deletions. The resulting alignment gaps are treated as unresolved characters that effectively do not contribute to the likelihood [103]. Deletions (and insertions) in coding regions arguably introduce changes that may disrupt protein function [104]. However, they do not register as an increase in the rate of protein evolution. Because deletions are not reversible events [105], they cannot be straightforwardly embedded into most common frameworks. Meanwhile, a high rate of codon deletion could be interpreted as evidence of a propensity to fix slightly deleterious changes.

Is there other evidence of potentially deleterious changes being fixed in Myxozoa? A preference for arginine over lysine and for valine over isoleucine has been reported in vertebrates with larger effective populations (*N*_*e*_), possibly due to effects on protein stability [106]. Myxozoan sequences are indeed relatively higher in lysine and isoleucine, but their usage also correlates with GC content (Figure S9-S10) [107]. The association between *N*_*e*_ and GC, mediated by GC-biased gene conversion, further complicates the interpretation of the observation [108, 109]. An assessment of fold stability could hence be informative. Frequent intron retention has also been reported in Myxozoa, and could, in principle, be consistent with disrupted regulation of splicing due to the accumulation of deleterious mutations [4, 110, 111].

Genome structure may also provide hints about the balance of forces shaping myxozoan evolution. At first glance, the high degree of gene order conservation seen in *Kudoa* seems at odds with the dramatic shifts in genome composition that occurred in the myxozoan ancestor, as well as the notion that myxozoans are inherently fast-evolving. We might not expect gene order to be tightly constrained to ensure coordinated expression in a genome where erroneous transcripts are common (it is currently not known how gene expression varies across the genome). On the other hand, conservation need not result from selection to maintain gene order. Interestingly, the highly reduced bacterial endosymbiont *Buchnera* also shows little evidence of rearrangement [112–114]. The genomes of bacterial endosymbionts with established host relationships are reported to be stable, following an initial period of rapid change [115, 116]. In *Buchnera*, it has been hypothesised that static gene order is driven by a lack of recombination, either due to limited opportunity or loss of the required machinery [112]. Although myxozoans are thought to undergo meiosis in the annelid-infecting stage, it is not clear to what extent outcrossing occurs [8]. Due to limited read coverage, it was not possible to reliably estimate heterozygosity for *Kudoa*, though sequence diversity appears low (see Figure S15). Additional data would therefore be required to quantify the contribution of recombination to observed sequence variation. A low effective recombination rate and low heterozygosity could, meanwhile, both contribute to the observed reduction in GC relative to freeliving cnidarians [117]. It is also interesting to note that there is evidence of high rates of rearrangement in myxozoan mitochondrial genomes [99].

A reduced genome could be read as evidence of a well-adapted organism that has discarded components that are no longer required. However, it is necessary to distinguish between adaptive streamlining and deterioration. Gene losses might also occur when the mutation rate exceeds the fitness advantage of maintaining them, which will result in the removal of less essential sequences. Increased drift due to bottlenecks could have a similar outcome (see [12]). Establishing the relative impacts of these processes on genome reduction is challenging, in part because we can only speculate about fitness effects associated with individual losses. In addition, it is not obvious if parasites become smaller before their genomes do [93]. It is, therefore, interesting to consider the degree of contraction across different sequence classes. Simulations in bacteria predict that increased mutation combined with reduced *N*_*e*_ leads to both coding and non-coding sequence loss. Meanwhile, more effective selection leads to more compact genomes that maintain their coding content while removing non-coding sequence [116].

Although it is not clear how well these predictions carry over to eukaryotes, the situation in *Kudoa* appears closer to the first scenario. Both *Kudoa* genomes examined in this work contain a relatively high fraction of repeats, the majority of which are unlikely to serve a function – particularly those showing substantial degradation. Repeat content is in fact higher than in some free-living cnidarians with considerably larger gene repertoires, such as *N. vectensis* (31%). Similar observations have been made in other myxozoans, as well as in rhizarian parasites [91, 118]. It has been suggested that the loss of cytosine methylation in *Myxosporea* could relate to a reduced need for small genomes to defend themselves against transposons [3]. Whether this is true or not, it is not known if genome contraction predates the loss of the components required for cytosine methylation. A reduced capacity to suppress expansion and purge existing elements could nevertheless result in an abundance of repeats. Neither *K. neothunni* nor *K. sp. trachurus* showed evidence of the methylation-related genes reported absent in other myxosporeans, based on a search of the proteome with HMM profiles provided by Kyger et al. [3].

This work demonstrates that sequencing data from infected host species can reveal genomes from under-sampled parasitic organisms. Many questions about how myxozoans evolve remain, however, given the limitations of relying primarily on coincidentally discovered genomes. Additional data that allow *N*_*e*_ and rates of outcrossing and mutation to be estimated would be especially useful. Quantifying biases in comparisons between fast-evolving sparsely sampled groups and more comprehensively sequenced, slower-evolving ones is also of interest. The circumstances where rate estimates are likely to be affected will have implications for assessing how selection shapes proteins in fast-diverging parasitic groups with limited genomic data. Beyond expanding efforts to obtain additional genomes, models that accommodate heterogeneity in amino acid preferences among sites could help address these issues in the future (see [101]).

## ACKNOWLEDGEMENTS

We would like to acknowledge our colleagues at the Tree of Life for many helpful conversations about cobiont assemblies and contributions to the underlying datasets. Particular thanks go to James Torrance for helping with the genome submissions. We also thank Sally Chang for providing access to *K. iwatai* protein sequences, Dorothée Huchon for useful pointers on myxozoans, and Nick Goldman and Nicola de Maio for helpful discussions.

This work was funded by Wellcome Trust Grants 206194 and 218328. For the purpose of Open Access, the author has applied a CC BY public copyright licence to any Author Accepted Manuscript version arising from this submission.

## DATA AVAILABILITY

The code for generating network embeddings and visualising Hi-C maps is available from https://github.com/CobiontID/HiC_network under an MIT license.

Both *Kudoa* assemblies described in this work have been submitted to ENA. Data associated with *K. neothunni* (jmKudNeot1) have been linked to Biosample SAMEA115884706. Data for *K. sp. trachurus* (jmKudSpea1) have been linked to Biosample SAMEA116001081. See https://github.com/CobiontID/Kudoa_genomes for an overview. Additional read sets generated for this work are available under run accessions ERR13510304, ERR13510305, and ERR13510306. The updated *Th. albacares* assembly, from which additional myxozoan sequence was removed, is available under the accession GCA_914725855.2.

