## Supplementary Material for "*Kudoa* genomes from contaminated hosts reveal extensive gene order conservation and rapid sequence evolution"

#### Identification of contaminant scaffolds

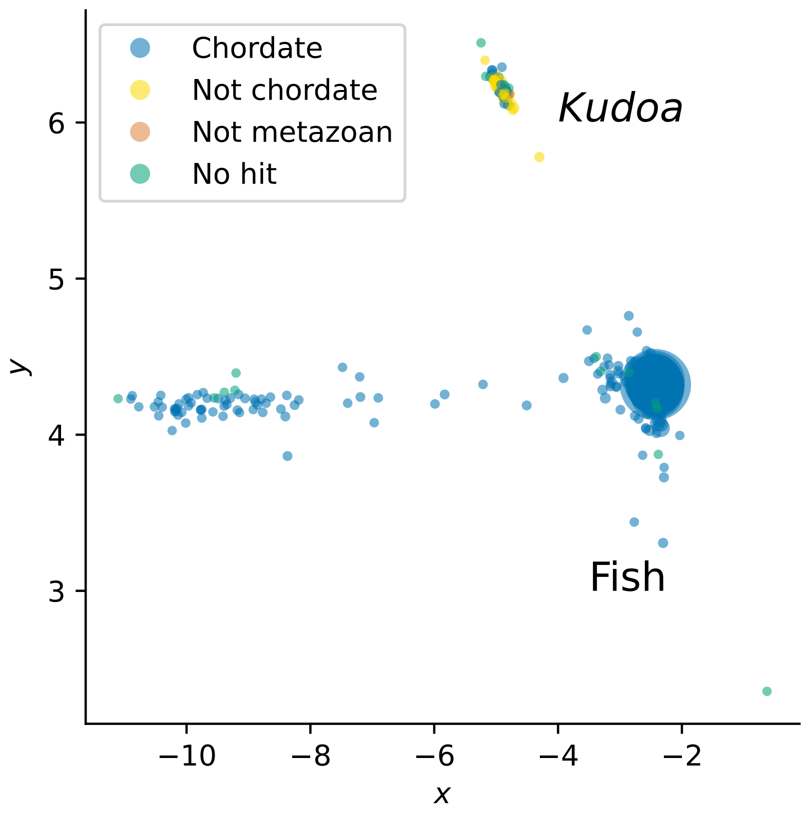


**Supplementary Figure 1.** 2D embeddings on scaffolds from initial *Tr. trachurus* assembly, coloured by buscoregions_phylum assignments from BTK

### Phylogenetic placement of marker sequences

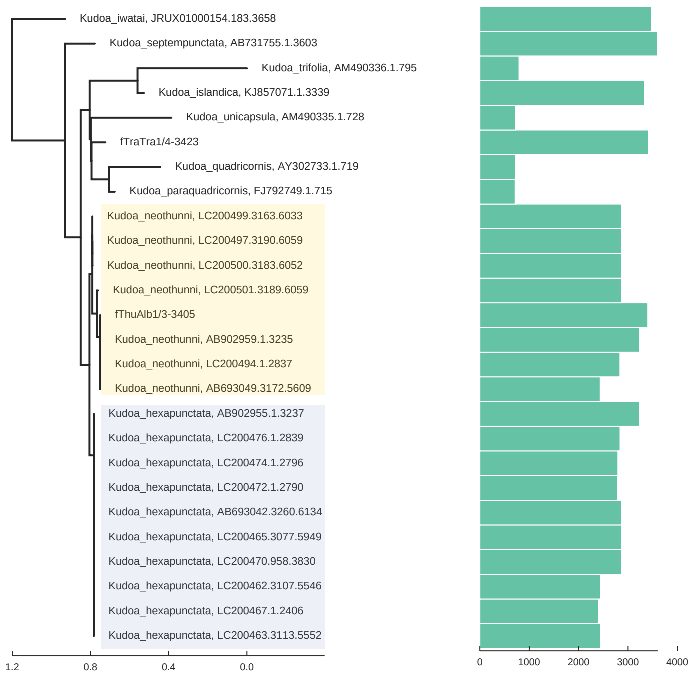


**Supplementary Figure 2.** Tree of *Kudoa* nLSU sequences from SILVA DB, aligned with Infernal. The phylogeny was generated in RAxML under the GTR+G/16A+G secondary structure model. The parasite found in *Th. albacares* is placed within *K. neothunni*, while the taxonomic assignment of the sequence found in *Tr. trachurus* is unclear. The bars in the column on the right indicate sequence length, and illustrate the lack of complete marker sequences for certain branches. Tip labels include accessions from which the 28S was extracted.

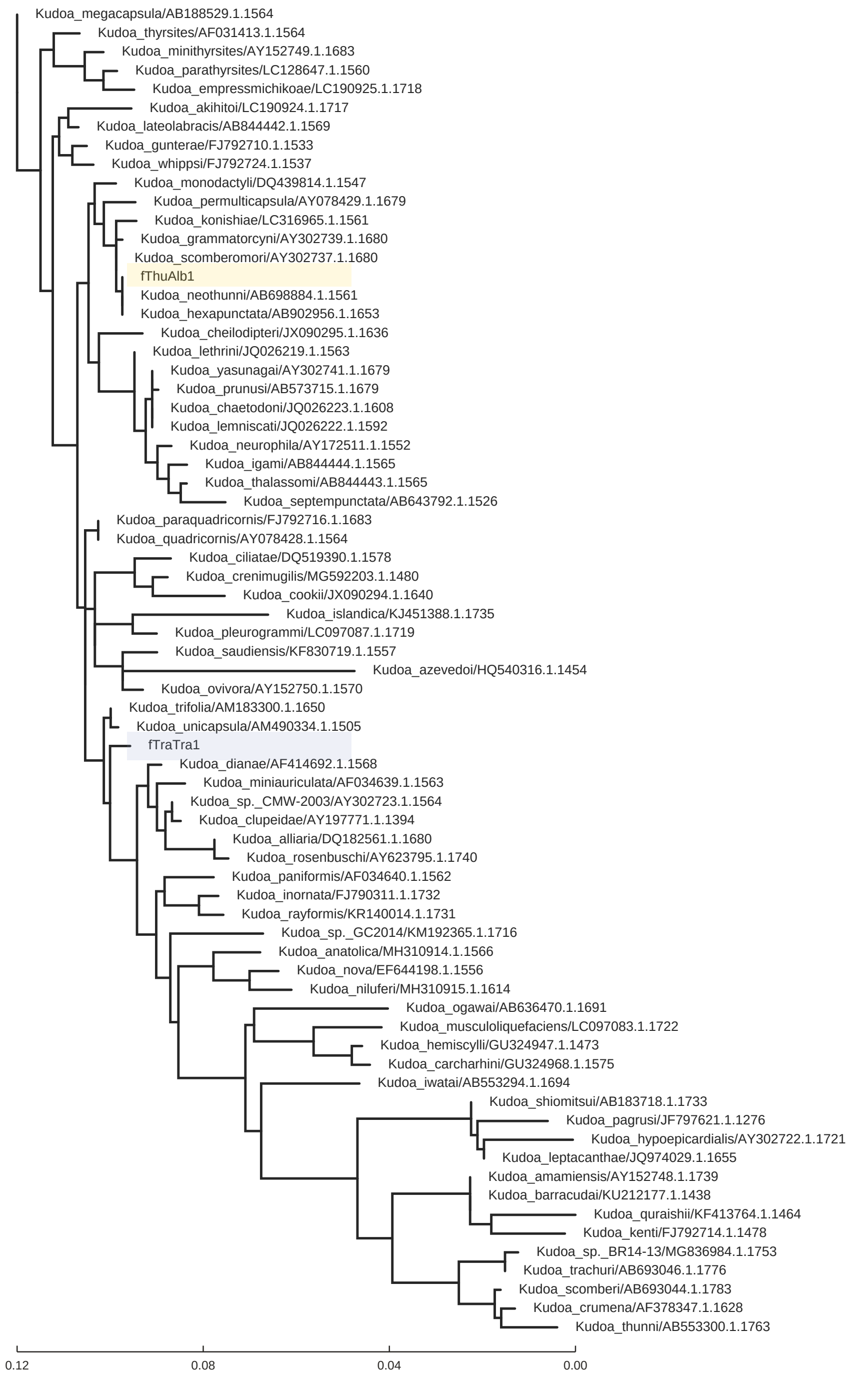


**Supplementary Figure 3.** Tree of Kudoa nSSU sequences from Silva DB, aligned with SSU-align, duplicate sequences removed, phylogeny inferred in RAxML under the GTR+G/16A+G secondary structure model. The sequences from the two fish assemblies are highlighted in colour. The *Kudoa* nSSU found in *Th. albacares* groups with *K. neothunni* and *K. hexapunctata*, but cannot be explicitly assigned due to insufficient divergence between the 18S sequences of the two species. Although the nSSU is better-represented in the database than the nLSU, the *Kudoa* 18S retrieved from *Tr. trachurus* cannot be assigned to a known species. For the purpose of visualisation, only one sequence per species is displayed, and truncated sequences <1000 bp were excluded. Tip labels include accessions.

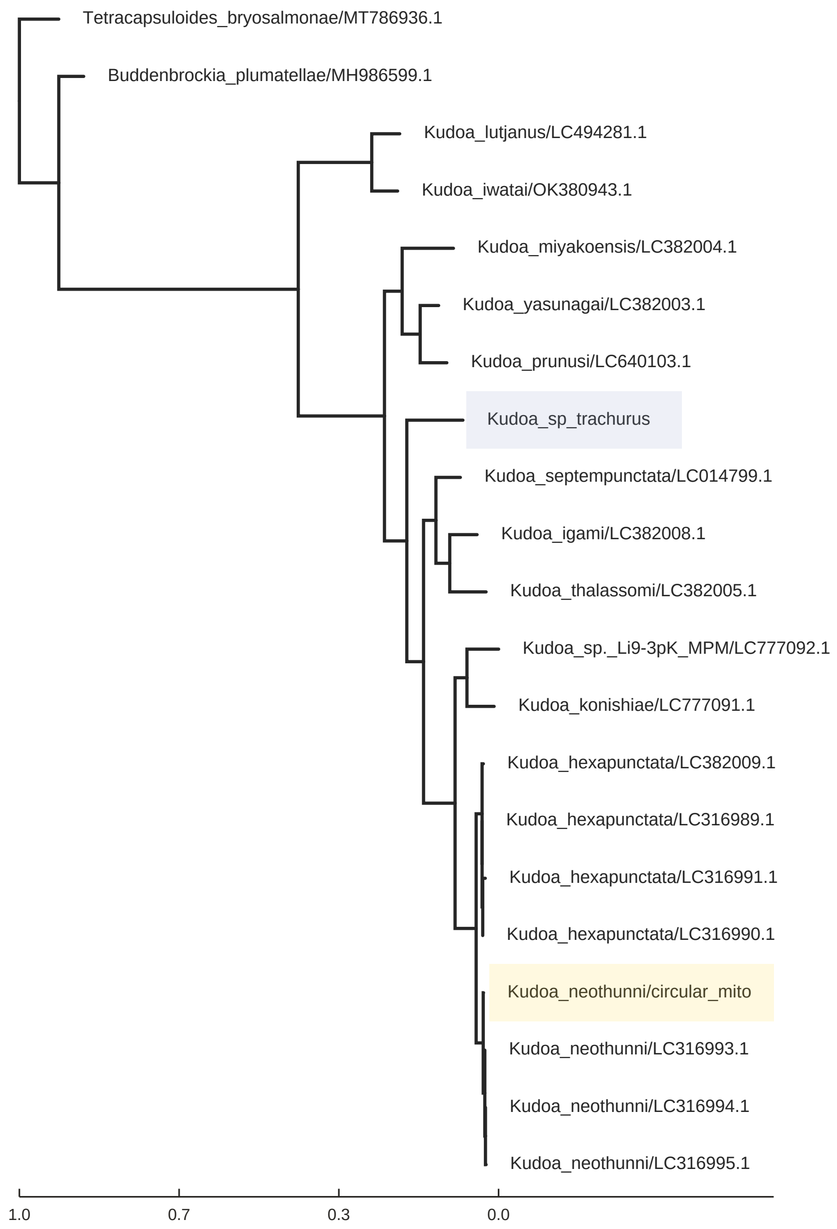


**Supplementary Figure 4.** Tree of *Kudoa* mitochondrial Cox1 sequences inferred under Kimura's two-parameter model in IQ-TREE2, confirming that the mitochondrial sequence retrieved from *Th. albacares* (highlighted in yellow) groups with *K. neothunni*, as is the case for the nLSU. The sequence from *Tr. trachurus* could, again, not be taxonomically assigned. The short distance between *K. neothunni* and *K. hexapunctata* prompted the choice of a nucleotide model.

### Hi-C network embeddings

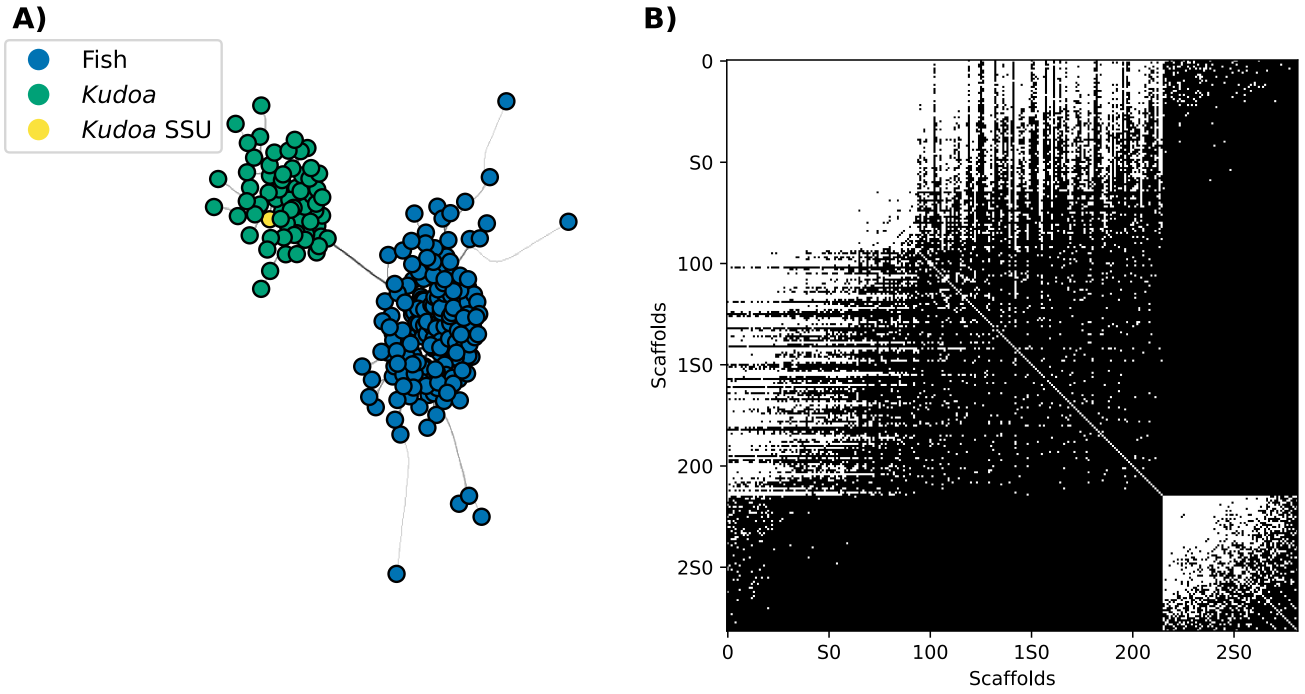


**Supplementary Figure 5.** Network analysis for the preliminary *Tr. trachurus* assembly. **A)** Displays the network produced from the scaffold adjacencies, with grey lines representing connections, and scaffolds containing the *Kudoa* nSSU marked yellow. Blue and yellow show the cluster labels assigned by node2vec. **B)** Shows the binary adjacency matrix for all contig pairs sorted by clusters, with *Kudoa* at the bottom right. White pixels indicate the presence of at least one connection. See Figure 2 for equivalent assignments for *Th. albacares*.

As noted in the methods, the node2vec algorithm performs biased random walks over the graph, which are then used to train a skipgram model. The parameters $p$ and $q$ affect how quickly the walk will tend to leave the local neighbourhood, which in turn affects whether nodes that are highly interconnected will be assigned similar embeddings, as opposed to having similar structural roles in the network. The former is usually achieved by setting $p$ higher and $q$ lower (e.g. $p = 1$, $q = 0.5$). In practice, identical label predictions were obtained under a range of parameter settings.

The presence of spurious connections between a subset of the fish and parasite scaffolds, illustrated in Figure 2, raises the question whether considering the number of connections between each pair of scaffolds would be useful. This is especially apparent in *Th. albacares*, and is visible in the layout of the NetworkX graph, which places a subset of the fish scaffolds closer to the parasite scaffolds. This is also apparent upon inspecting a 2D representation of the node2vec embeddings, but a clear affinity nevertheless remains between the fish scaffolds. When all off-diagonal matrix elements are included in the graph, effectively weighting the edges, the pattern disappears. However, this would result in the biased random walk tending to visit large scaffolds with many connections more frequently. It also effectively overrides the $p$ and $q$ parameters. Filtering out multi-mapping Hi-C reads before constructing the adjacency matrix may therefore be more appropriate.

### Read distribution and coverage

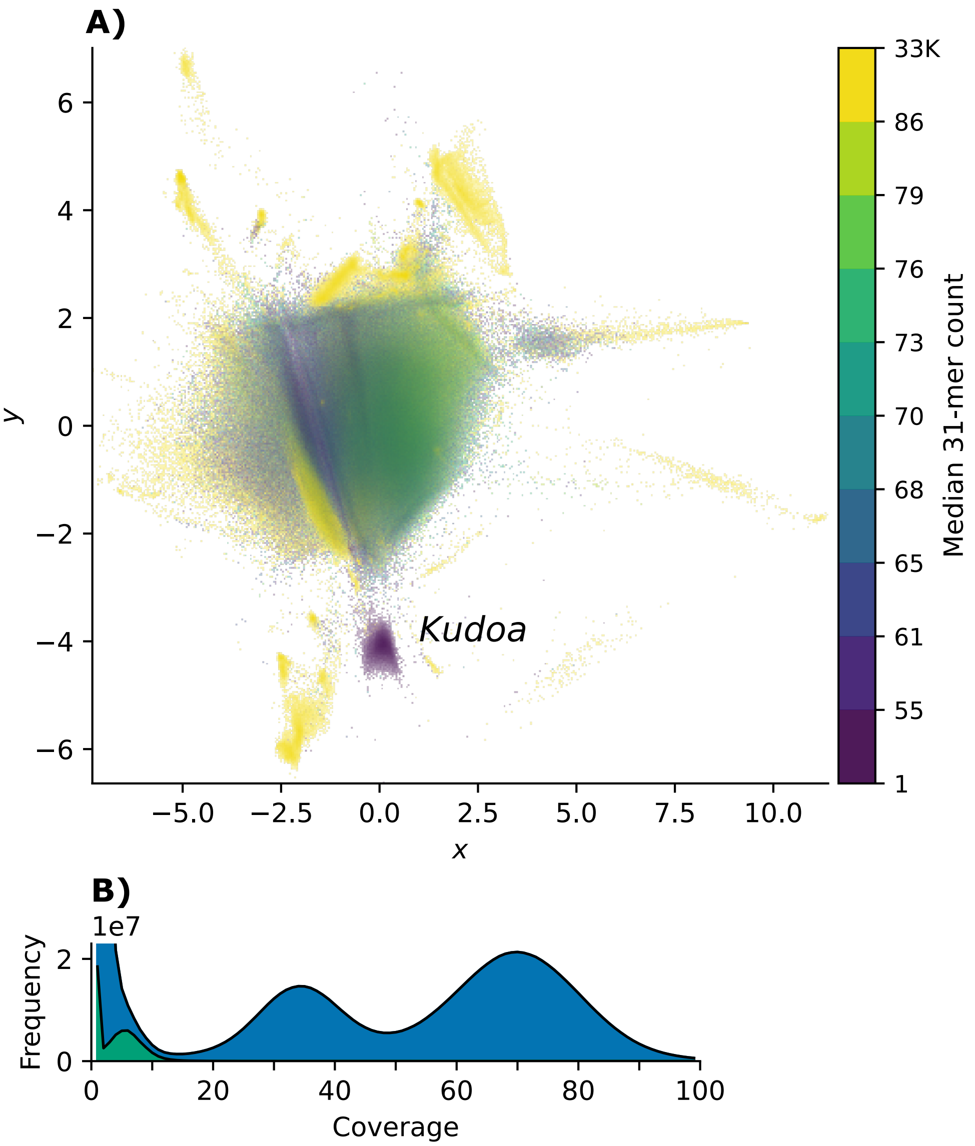


**Supplementary Figure 6.** Coverage of Kudoa reads in *Th. albacares.* **A)** shows 2D VAE embeddings of all reads from *Th. albacares*, coloured by k-mer coverage (k = 31). **B)** shows the k-mer distribution (k = 31) of the full read set (upper curve, shaded blue), with the distribution of the reads in the putative *Kudoa* cluster superimposed (lower curve, shaded green).

### Data sources

| | Class | Genome Accession | Other Source |
|---|---|---|---|
| *Acropora millepora* | *Anthozoa* | GCA_013753865.1 | – |
| *Actinia tenebrosa* | *Anthozoa* | GCA_009602425.1 | – |
| *Blastomussa wellsi* | *Anthozoa* | GCA_947652115.1 | – |
| *Ceratonova shasta* | *Myxozoa* | GCA_918559645.1 | https://doi.org/10.5061/dryad.tx95x69tt |
| *Exaiptasia diaphana* | *Anthozoa* | GCA_001417965.1 | – |
| *Galaxea fascicularis* | *Anthozo*a | GCA_948470475.1 | – |
| *Henneguya salminicola* | *Myxozoa* | GCA_009887335.1 | – |
| *Hydractinia symbiolongicarpus* | *Hydrozoa* | GCA_029227915.2 | – |
| *Hydra vulgaris* | *Hydrozoa* | GCA_022113875.1 | – |
| *Kudoa iwatai* | *Myxozoa* | GCA_001407235.2 | – |
| *Lumbricus rubellus* (outgroup) | *Clitellata* | GCA_945859605.1 | – |
| *Montipora capitata* | *Anthozoa* | GCA_949126865.1 | – |
| *Myxobolus honghuensis* | *Myxozoa* | GCA_022478855.1 | https://doi.org/10.7910/DVN/INLEPM |
| *Myxobolus squamalis* | *Myxozoa* | GCA_010108815.2 | – |
| *Nematostella vectensis* | *Anthozoa* | GCA_932526225.1 | – |
| *Orbicella faveolata* | *Anthozoa* | GCA_002042975.1 | – |
| *Pocillopora damicornis* | *Anthozoa* | GCA_003704095.1 | – |
| *Polypodium hydriforme* | *Polypodiozoa* | – | PRJNA251648 (transcriptome) |
| *Porites lutea* | *Anthozoa* | GCA_958299795.1 | – |
| *Ricordea florida* | *Anthozoa* | GCA_949710005.1 | – |
| *Sphaerospora molnari* | *Myxozoa* | – | https://doi.org/10.5061/dryad.j3tx95x9c |
| *Stylopora pistillata* | *Anthozoa* | GCA_002571385.2 | – |
| *Tetracapsuloides bryosalmonae* | *Myxozoa* | – | https://doi.org/10.6084/m9.figshare.13302746.v1 |
| *Thelohanellus kitaei* | *Myxozoa* | GCA_000827895.1 | – |

**Supplementary Table 1.** Sources for publicly available genomic or transcriptomic data.

### Annotation stats

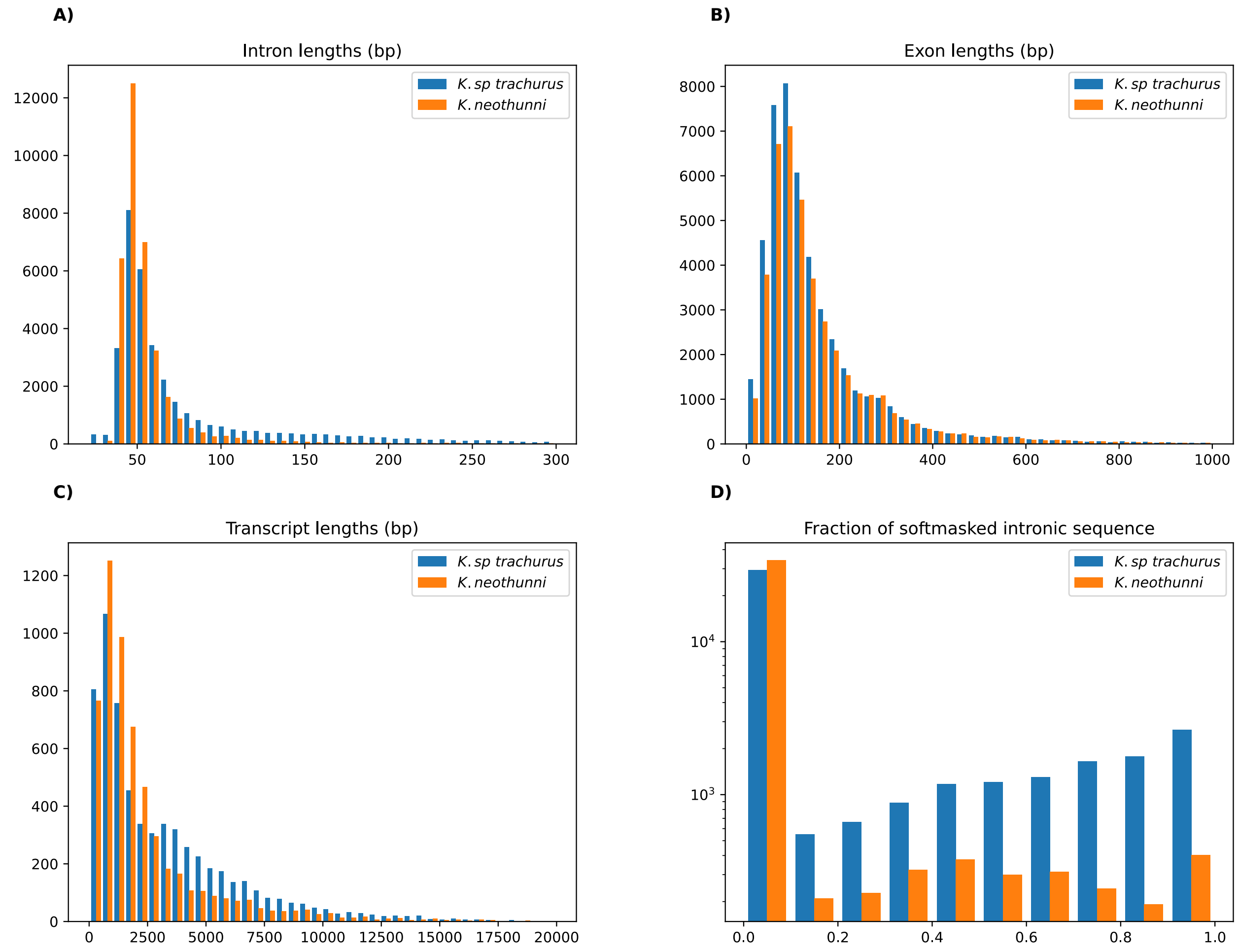


**Supplementary Figure 7.** Feature length distribution. While the distribution of exon sizes is broadly similar between both species, K. *sp. trachurus* appears to contain more long introns than *K. neothunni*. A similar pattern is seen for transcript length, perhaps explained by a larger fraction of repeats in *K. sp. trachurus*. AUGUSTUS defines transcript length as the distance between the start and stop codon. Therefore, each predicted transcript is unspliced and includes both exons and introns.

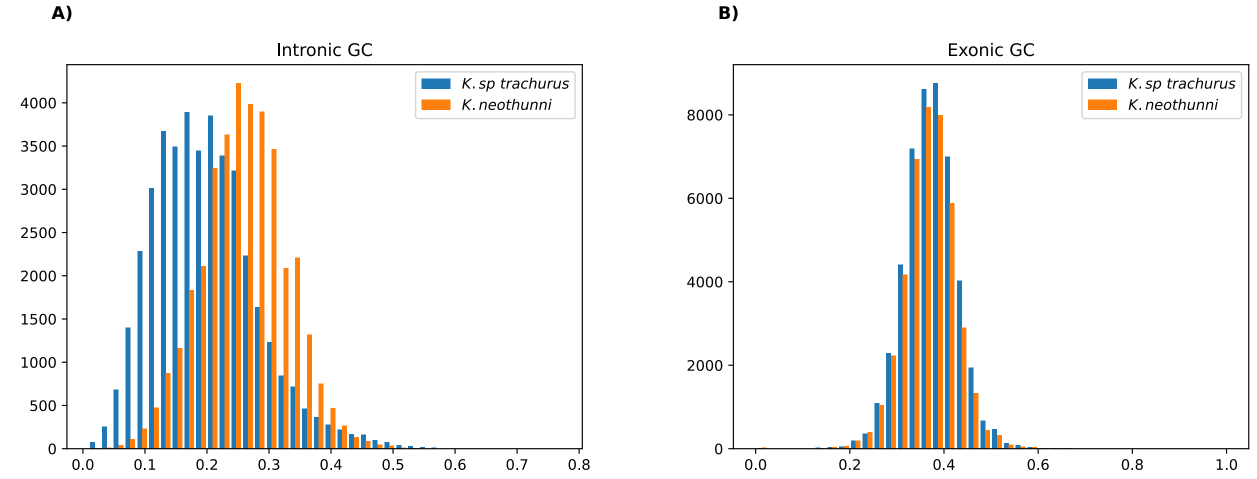


**Supplementary Figure 8.** GC content in *K. sp. trachurus* is lower than in *K. neothunni* in introns, but similar in exons.

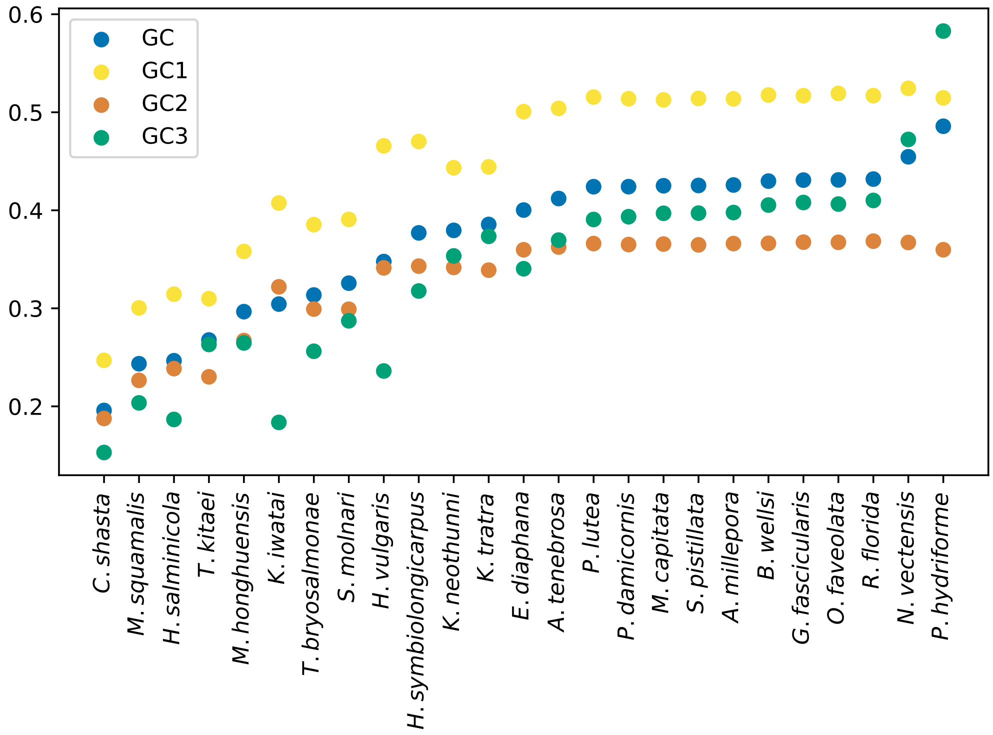


**Supplementary Figure 9.** GC content for each cnidarian species, for all sites in coding sequences and individual codon positions, based on the dataset used to examine codon substitution patterns. Corals show relatively similar composition, in line with shorter evolutionary distances, while myxozoans GC tends to be lower and more variable between species.

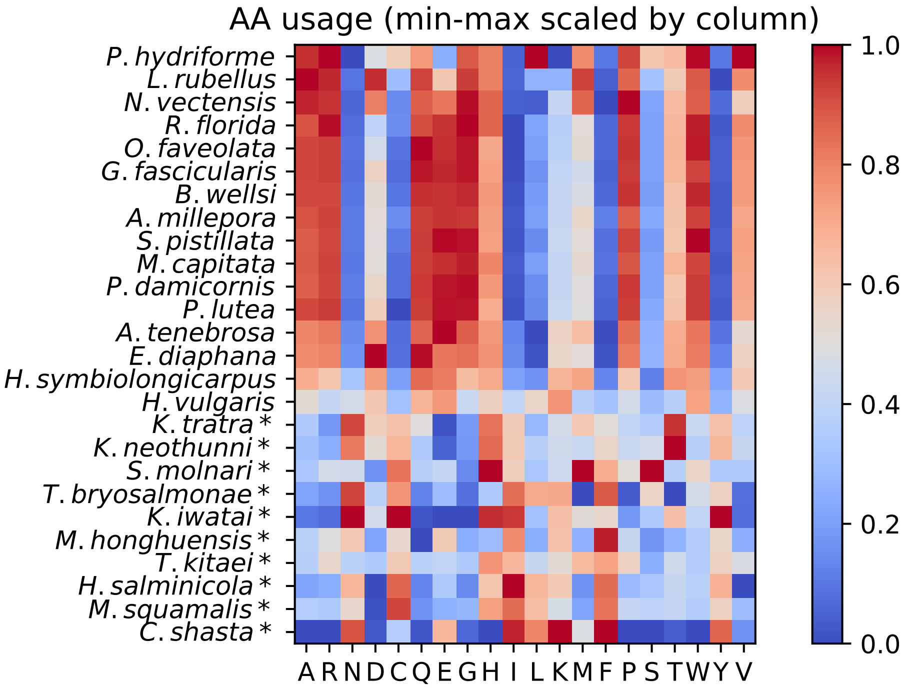


**Supplementary Figure 10.** Amino acid usage across the set of orthologs used to infer substitution rates, as above. The fraction for each amino acid is min-max scaled to highlight species with relatively higher usage. Myxozoan species are marked with an asterisk.

### Repeat distribution

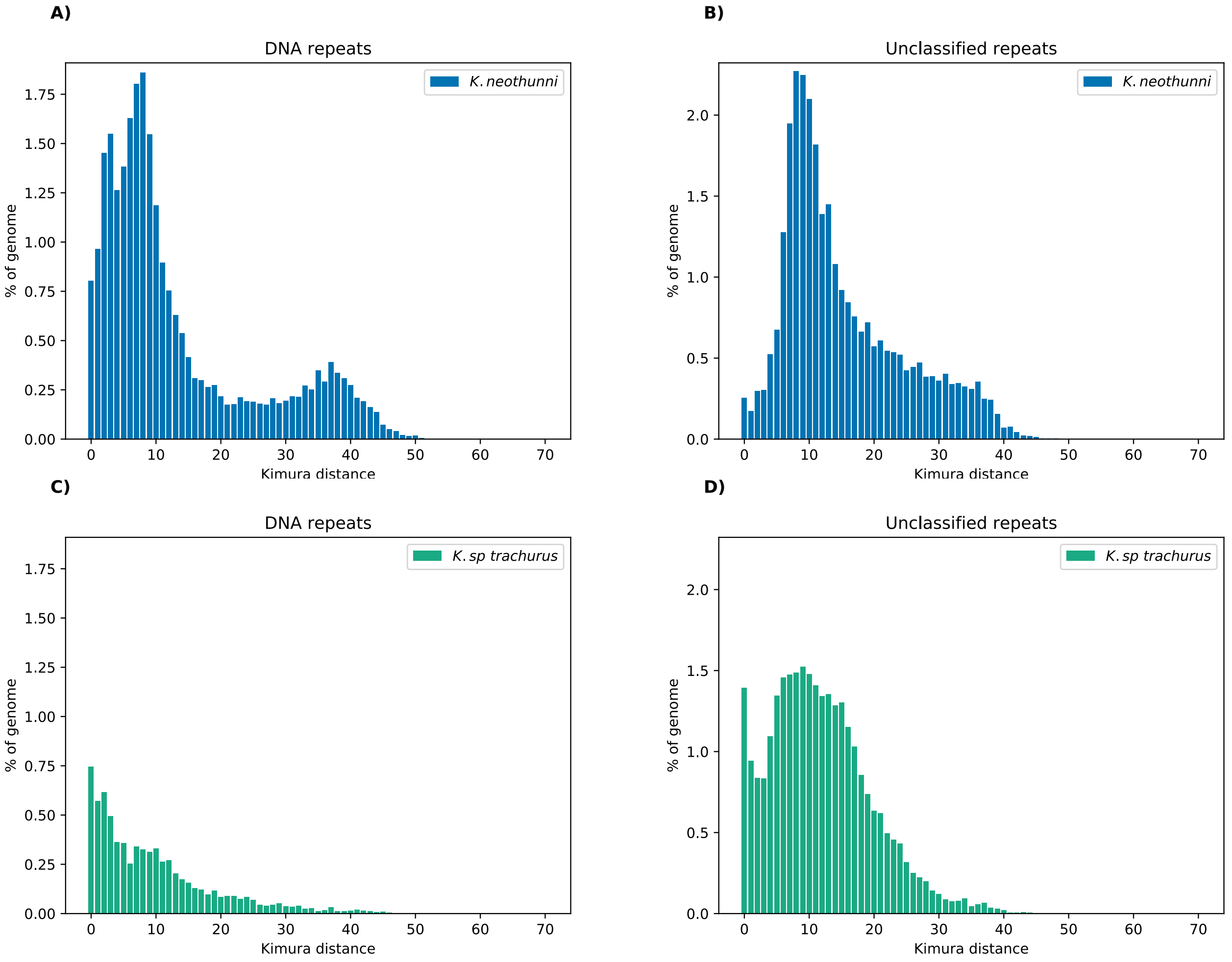


**Supplementary Figure 11.** Summary of repeat content and divergence from consensus sequences, as estimated by Earl Grey for *K. neothunni* and *K. sp. trachurus*. Only sequences marked as “Unclassified” or “DNA” repeats are shown, as these account for the vast majority of elements. The distribution for *K. neothunni* implies a larger fraction of older, more diverged insertions.

### Synteny

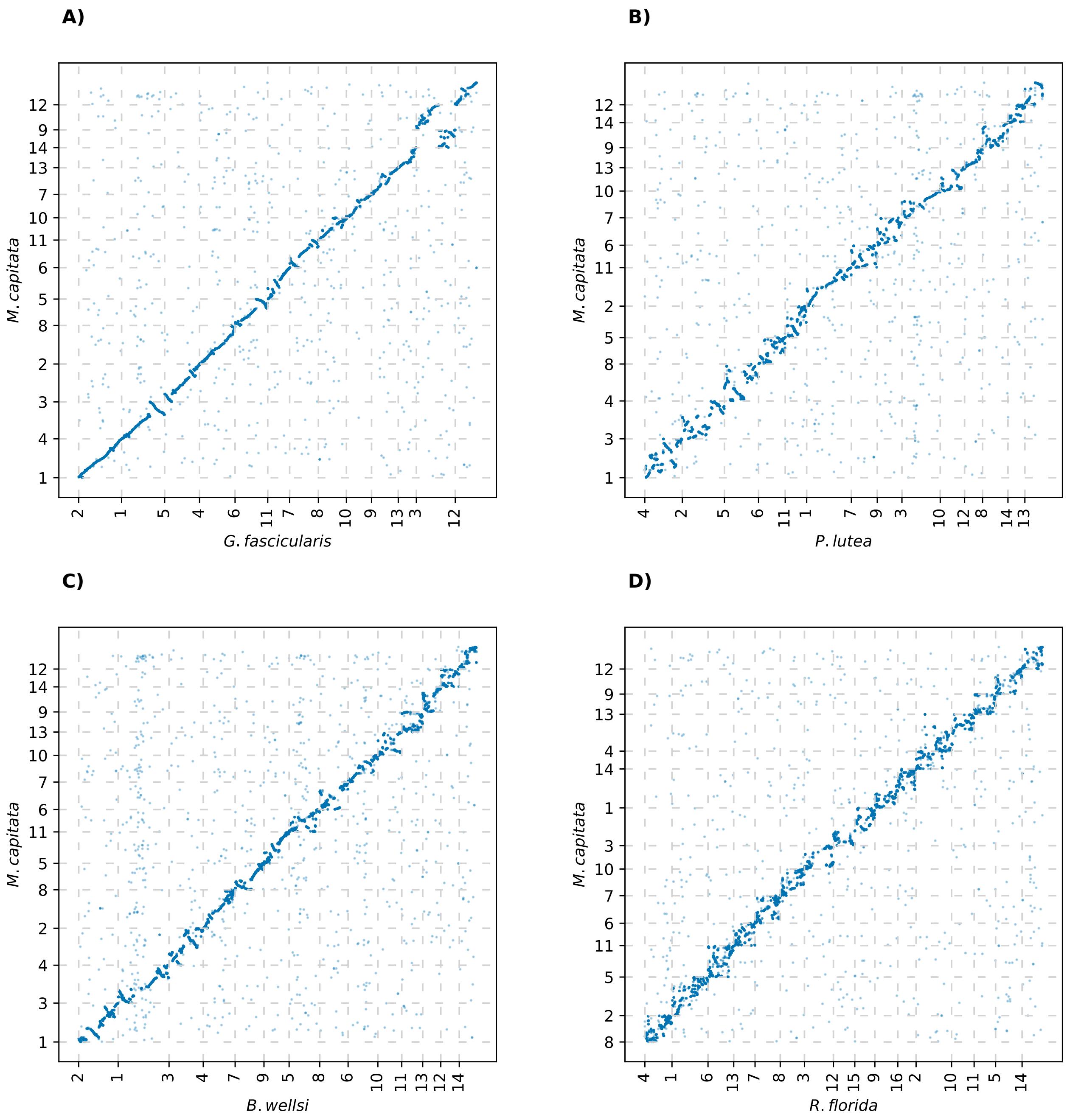


**Supplementary Figure 12.** Synteny conservation for orthologs between *M. capitata* and four other coral species. The panels are sorted by inferred amino acid divergence in descending order ($t = 0.22, 0.28, 0.32, 0.44$). Microsynteny degrades with increased evolutionary distance, though inferred divergence appears to be an imperfect predictor. Divergence between *M. capitata* and *B. wellsi* is approximately on par with *K. neothunni* and *K. sp. trachurus*, which are shown in Figure 7. Distances are based on the PMSF tree in Figure 6.

### Substitution rates

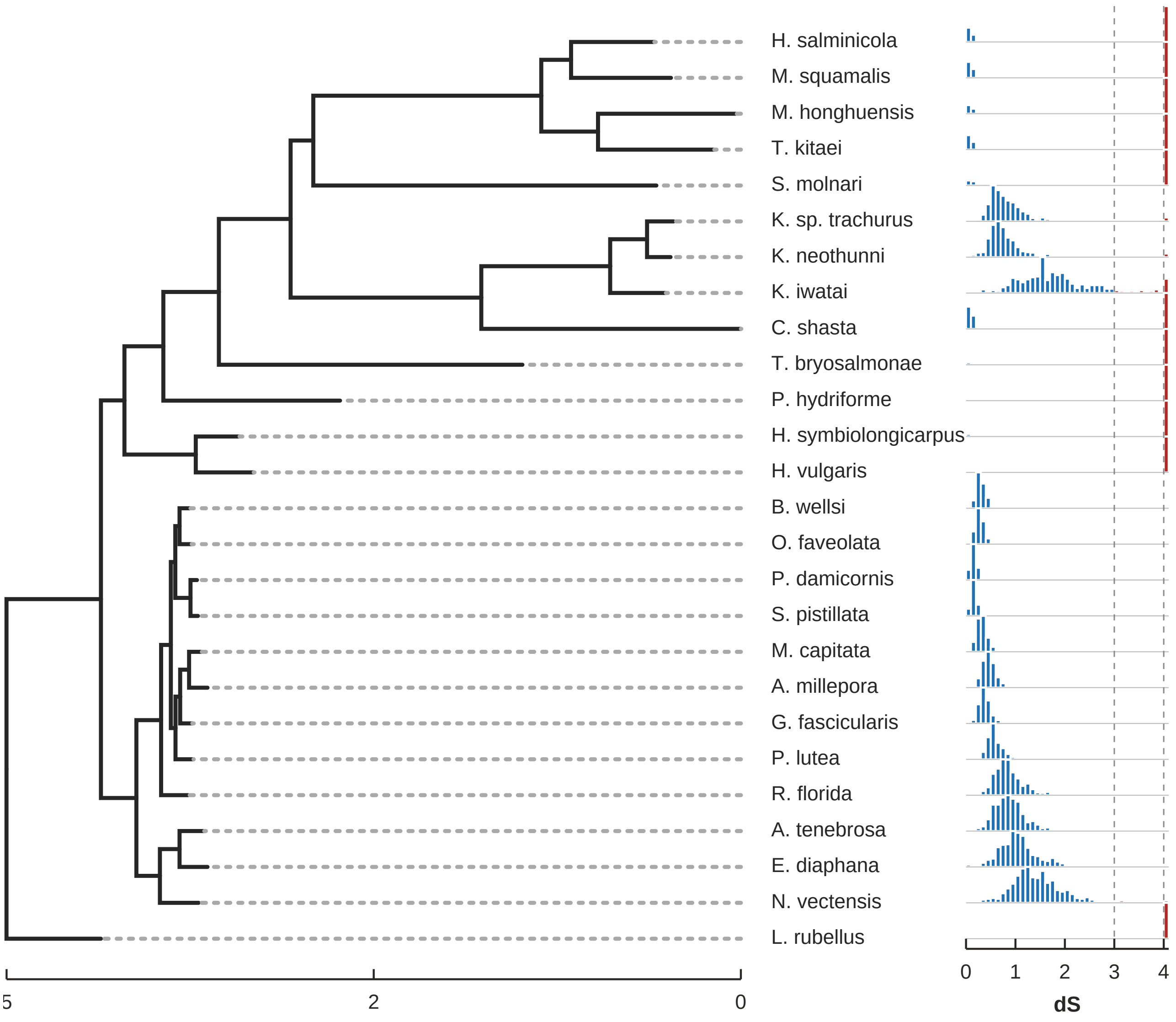


**Supplementary Figure 13.** A PMSF tree for Cnidaria annotated with information about synonymous divergence highlights groups likely to yield unreliable parameter estimates. Branch-specific values were obtained with codeml for all alignments in the "extended" dataset under M2 as described in the Methods. The histograms indicate the distribution of $dS$ for each tip, with the last bin containing all values greater than 4. Values below 3 are shown in blue, and values above or equal to 3 are shown in red. A suggested rule of thumb is to treat alignments where $dS > 3$ with caution [79].

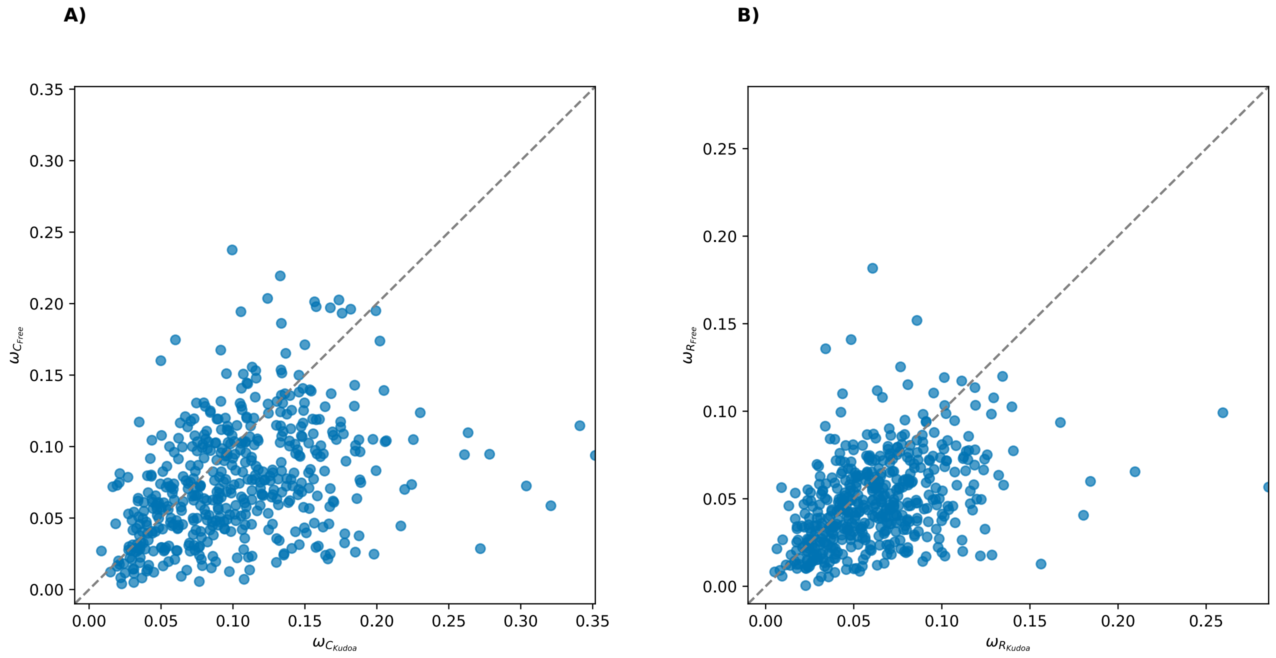


**Supplementary Figure 14.** Comparison of estimates of $\omega$ for *Kudoa* versus free-living species, considering **A)** conservative amino acid changes ($\omega_C$) and **B)** radical amino acid changes ($\omega_R$) separately. Estimates are overall consistent with slightly stronger selection against radical substitutions, but the difference between clades for the more constrained radical substitutions remains modest. Interestingly, a preference for conservative over radical changes is not generally consistently seen for low-$N_e$ species [80], but the removal of less conserved residues in this analysis may affect the results. Note: If $\omega$ is underestimated due to site-specific amino acid preferences being disregarded, this is also expected to affect $\omega_C$ and $\omega_R$.

### Genetic variation in *Kudoa*

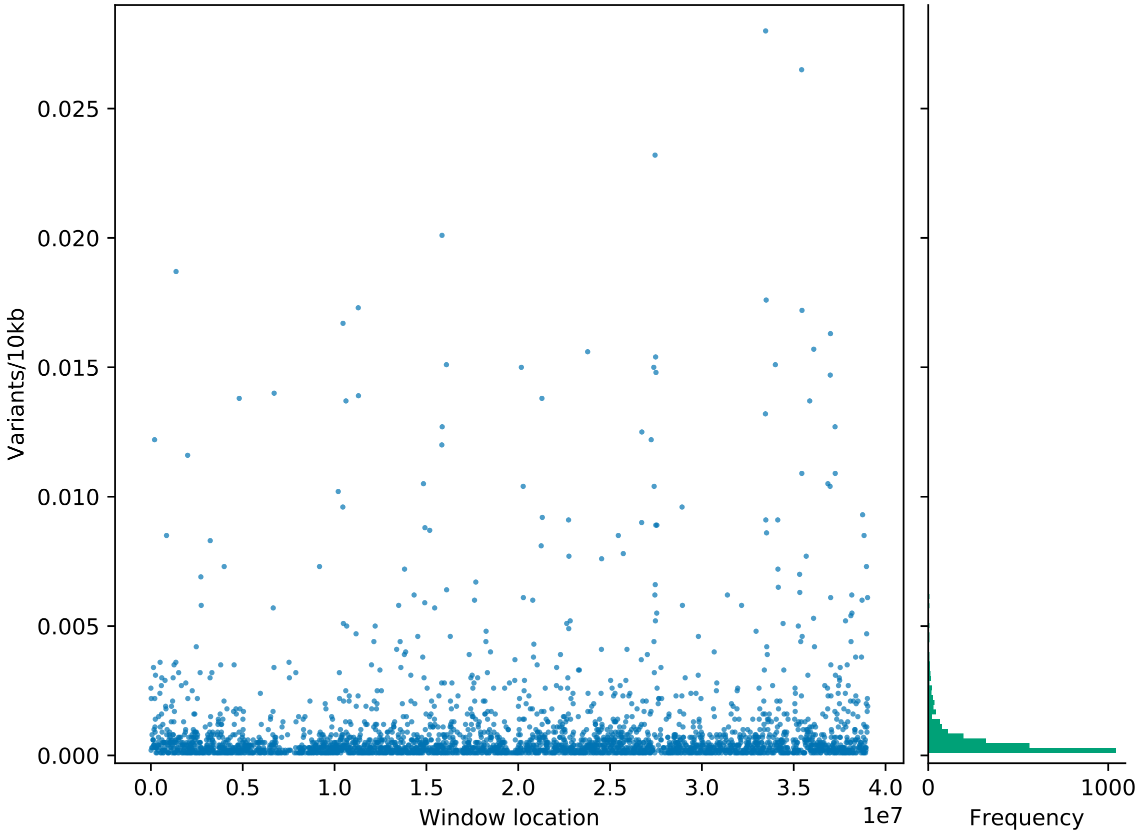


**Supplementary Figure 15.** Variants per 10 kb window for scaffold 4 in *K. neothunni*. Variants were called with DeepVariant [119]. Due to low coverage, a subset of windows contained no callable sites. Reliable estimates of $\pi$ per window could therefore not be obtained. Windows without calls are not displayed. For windows with calls, the median number of variants per 10 kb is 4.

### Telomeric repeats in *K. neothunni*

Two scaffolds in the uncurated assembly contained 96 and 88 instances of the motif "AAACCTAAACCT" within the first 1000 base pair window, respectively, and were each derived from a single contig from the primary meta-assembly. An additional contig that was placed in the alternate assembly, contained 60 instances of the motif. In the curated assembly, telomeric repeats are retained at one end of Chromosome 2. The number of reads containing the telomeric motif (n = 49) suggests that the remaining ends failed to assemble, which is not unexpected given the repetitive nature of the subtelomeric sequences.
